# Quantitative *in vivo* analyses reveal a complex pharmacogenomic landscape in lung adenocarcinoma

**DOI:** 10.1101/2020.01.28.923912

**Authors:** Chuan Li, Wen-Yang Lin, Hira Rizvi, Hongchen Cai, Christopher D. McFarland, Zoë N. Rogers, Maryam Yousefi, Ian P. Winters, Charles M. Rudin, Dmitri A. Petrov, Monte M. Winslow

## Abstract

The lack of knowledge about the relationship between tumor genotypes and therapeutic responses remains one of the most important gaps in enabling the effective use of cancer therapies. Here, we couple a multiplexed and quantitative platform with robust statistical methods to enable pharmacogenomic mapping of lung cancer treatment responses *in vivo*. We uncover a surprisingly complex map of genotype-specific therapeutic responses, with over 20% of possible interactions showing significant resistance or sensitivity. We validate one of these interactions - the resistance of Keap1 mutant tumors to platinum therapy - using a large patient response dataset. Our results highlight the importance of understanding the genetic determinants of treatment responses in the development of precision therapies and define a strategy to identify such determinants.

## Main Text

Efforts over the past decade have generated many novel cancer therapies. However, the lack of understanding of the relationship between tumor genotype and therapeutic response remains a major challenge in translating these therapies into precision cancer treatments ^1^. While the genetic complexity of human cancer could in principle thwart therapeutic efforts that disregard biomarker-driven patient selection, it remains unclear the extent to which tumor suppressor gene alterations impact therapeutic responses ^2^. Precisely mapping the pharmacogenomic landscape should help design better clinical trials and select more effective therapies for individual patients.

The pharmacogenomic landscape of cancer drug responses has been investigated using cell lines, patient-derived xenografts (PDXs), and patients themselves ^3–6^. However, cell lines grown *in vitro* lack the appropriate *in vivo* environment and the genotypic heterogeneity of patient tumors. PDXs and human cell line transplantation models recapitulate some aspects of *in vivo* growth, but growth factor/receptor incompatibility, growth in non-orthotopic sites, and the obligate absence of the adaptive immune system compromise these approaches ^7–9^. Furthermore, human tumor-derived systems almost invariably have large numbers of mutations and genomic rearrangements, and thus even large-scale analyses often lack the statistical power to glean cause-and-effect relationships between individual genetic alterations and therapeutic responses^3,10^.

Genetically engineered mouse models of cancer uniquely enable the introduction of defined genetic alterations into adult somatic cells, which leads to the generation of autochthonous tumors ^11^. These tumors can recapitulate the genomic alterations, gene expression state, histopathology, and therapy-refractive nature of corresponding human cancers ^12, 13^. Despite the potential value of these models in pre-clinical translation studies, the breadth of their utility has been limited by the fact that they are neither readily scalable nor sufficiently quantitative ^14^.

To increase the scope and precision of *in vivo* cancer modeling and to assess tumor suppressor gene function in a multiplexed manner, we recently developed a method based on tumor-barcoding coupled with high-throughput barcode sequencing (Tuba-seq). This method integrates CRISPR/Cas9-based somatic genome engineering and molecular barcoding into well-established Cre/Lox-based genetically engineered mouse models of oncogenic Kras-driven lung cancer ^14^. The initiation of lung tumors with pools of barcoded Lenti-sgRNA/Cre viral vectors enables the generation of many tumors of different genotypes in parallel. All neoplastic cells within each clonal tumor have the same two-component barcode, where each sgID region identifies the sgRNA and the random barcode (BC) is unique to each tumor. Subsequent high-throughput sequencing of the sgID-BC region from bulk tumor-bearing lungs can quantify the number of neoplastic cells in each tumor of each genotype.

We initiated lung tumors in *Kras^LSL-G12D/+^;Rosa26^LSL-Tomato^;H11^LSL-Cas9^* (*KT;H11^LSL-Cas9^*) mice and control Cas9-negative *KT* mice with a pool of barcoded Lenti-sgRNA/Cre vectors targeting eleven putative tumor suppressors and four control vectors with inert sgRNAs (Lenti-sg*TS^Pool^*/Cre; **Fig. 1a**). Tumor suppressors were selected based on common occurrence in human lung adenocarcinomas and previously suggested roles in oncogenesis ^15^. 18 weeks after tumor initiation, the sgID-BC region from each bulk tumor-bearing lung was PCR amplified and sequenced to quantify tumor sizes (**Fig. 1a**).

**Figure 1.**
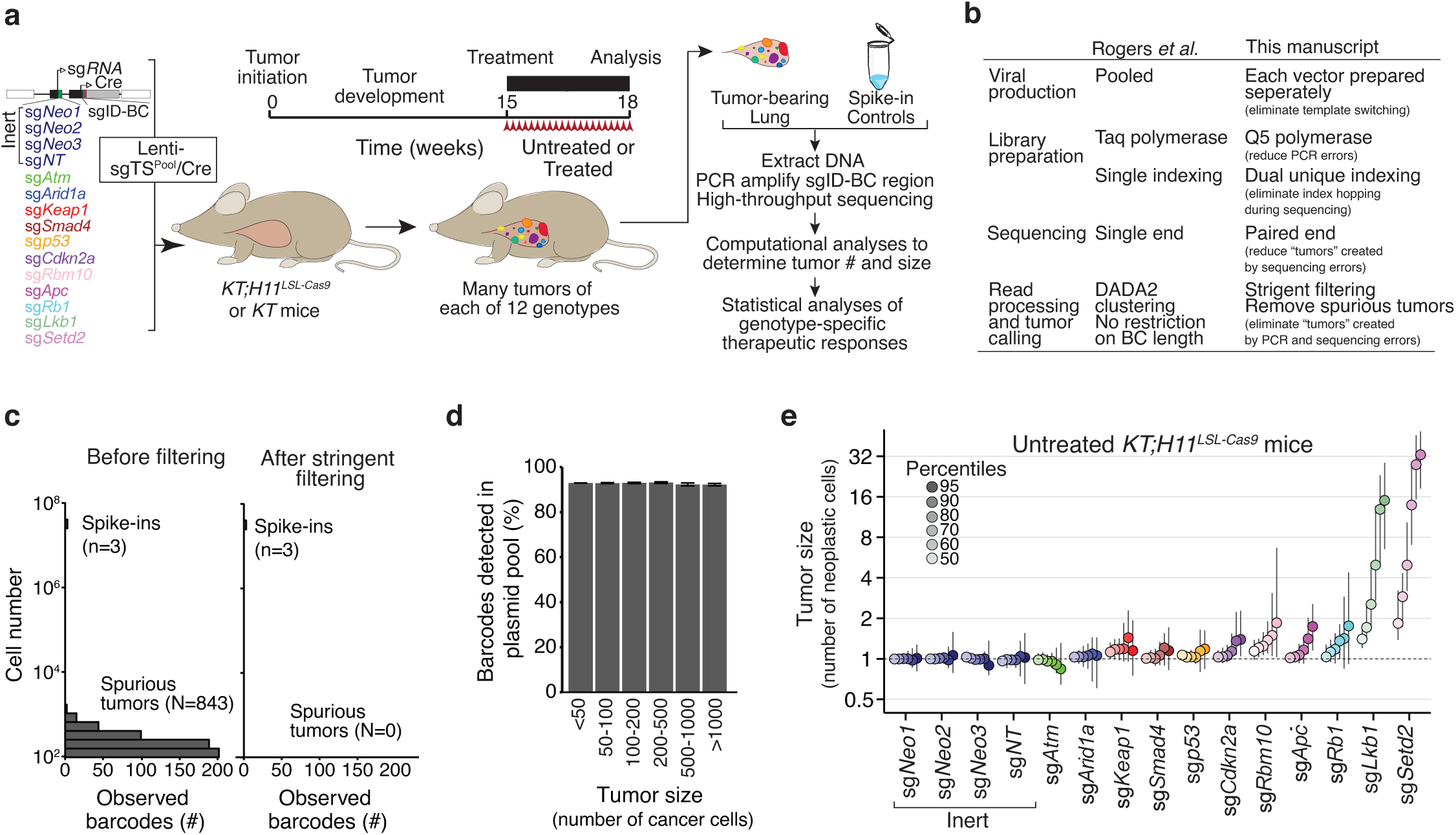
Optimization of tumor-barcoding coupled with high-throughput barcode sequencing (Tuba-seq) for the analysis of genotype-specific therapy responses (GSTRs) *in vivo*. **a.** Overview of Tuba-seq pipeline to uncover GSTRs. The Lenti-TS^Pool^/Cre viral pool contains barcoded vectors with sgRNAs targeting 11 putative tumor suppressors that are frequently mutated in human lung adenocarcinoma. Tumors are initiated in either *Kras^LSL-G12D/+^;R26^LSL-Tom^ (KT) or Kras^LSL-G12D/+^;R26^LSL-Tom^;H11^LSL-Cas9^ (KT;H11^LSL-Cas9^)* mice. Following tumor development, mice are treated with therapies, and barcode sequencing libraries are prepared from each tumor-bearing lung. **b.** Comparison of our current pipeline with our previous Tuba-seq pipeline (Rogers *et al.* 2017). **c.** Stringent filtering effectively eliminated spurious tumors. Analysis of the barcodes associated with the sgID specific for the Spike-in control cells (3 cell lines with a defined sgID-BC added at 5×10^5^ cell/sample as the benchmark) enables identification of recurrent barcode reads generated from sequencing and other errors (Spurious tumors). Data is from a typical lane of 22 multiplexed Tuba-seq libraries from *KT;H11^LSL-Cas9^* mice with Lenti-TS^Pool^/Cre initiated tumors. **d.** The percent of tumors with barcodes validated within the lentiviral plasmid pool is constant across tumor sizes. As sequencing and other processing errors are most likely to create small spurious tumors, this finding suggests that tumors detected by Tuba-seq represent real clonal expansions of barcoded cells. **e.** The relative size of tumors of each genotype in *KT;H11^LSL-Cas9^* mice 18 weeks after tumor initiation with Lenti-sgTS^Pool^/Cre. The relative sizes of tumors at the indicated percentiles were calculated from the tumor size distribution of all tumors in 5 mice. Error bars show 95% confidence intervals.

We optimized our Tuba-seq experimental protocols and analysis pipeline to eliminate sgRNA-sgID/barcode uncoupling due to lentiviral template switching and to minimize PCR, sequencing, and clustering errors (**Fig. 1b**, **Supplementary Fig. 1a,** and **Methods**) ^15^. Our new streamlined pipeline essentially eliminated the impact of read errors, as assessed by multiple metrics, including the remaining spurious tumors from spike-in barcodes with known sequences and correspondence of tumor barcodes with those from the lentiviral plasmid pool (**Fig. 1c, d**, **Supplementary Fig. 1b**). Quantification of the impact of tumor suppressor gene inactivation on tumor growth in *KT;H11^LSL-Cas9^* mice using our optimized method uncovered effects that were generally consistent with our previous analyses but with greater magnitudes of tumor suppression (**Fig. 1e**; **Supplementary Fig. 1c, d** and **2**, sign test for differences in magnitudes, *P* = 0.001) ^14^. Consistent with the robustness of our methods, analysis of the *KT* mice with Lenti-sg*TS^Pool^*/Cre-initiated tumor revealed no false-positive tumor suppressive effects (**Supplementary Fig. 1d, e**).

**Figure 2.**
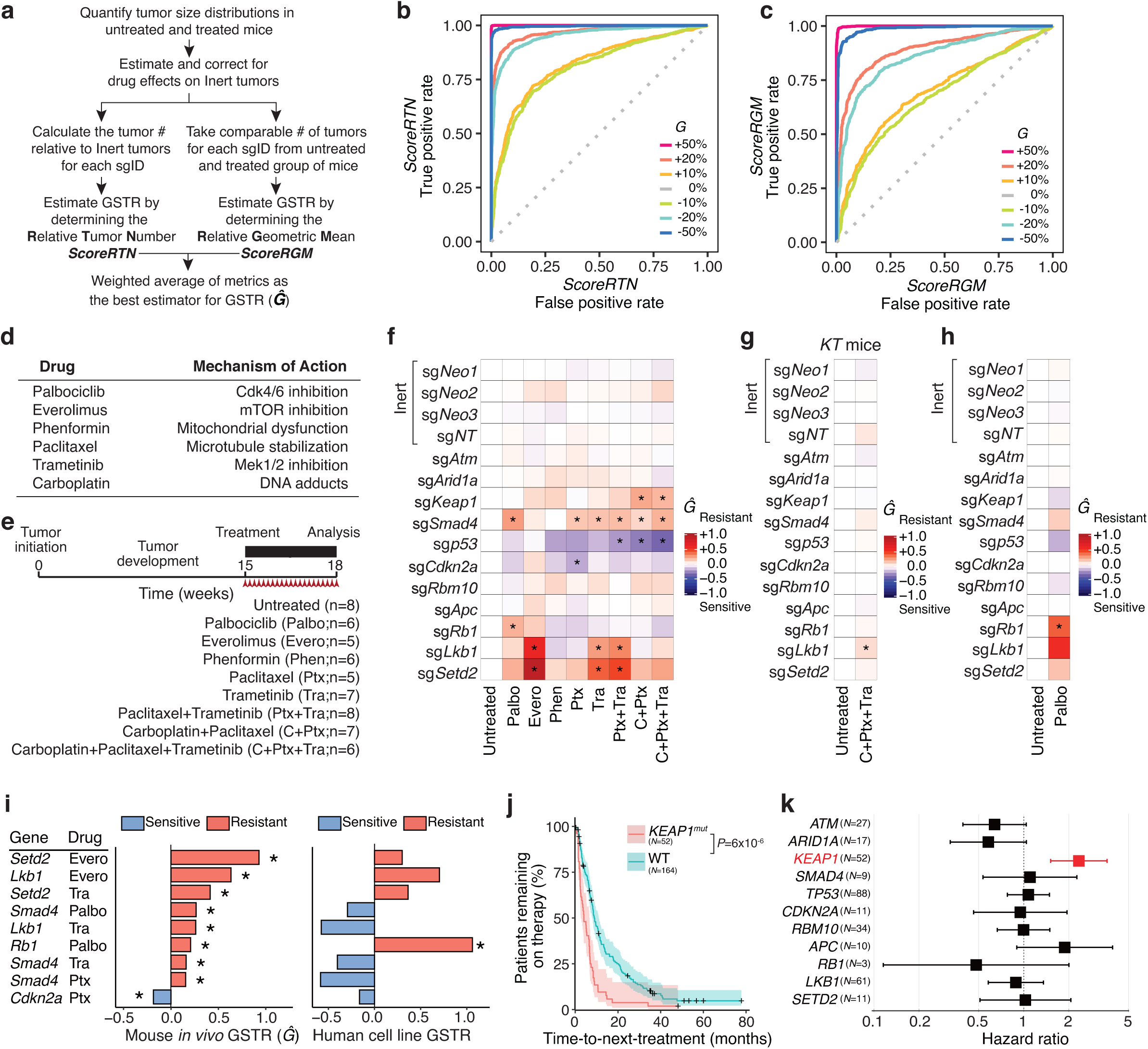
Tuba-seq quantifies genotype-specific therapeutic responses (GSTR) for multiple therapies. **a.** Data analysis pipeline to identify GSTR by comparing the relative tumor number (*ScoreRTN*) and relative geometric mean (*ScoreRGM*) between tumors containing a tumor suppressor targeting sgRNA and Inert tumors in the untreated and treated mice. **b.** The sensitivity and specificity of *ScoreRTN* estimated from simulations of preassigned drug effect (*S*=0.5) and GSTR (various *G*). There is no genotype-specific response when *G*=0. *G* of −20% means the tumors with the sgRNA were reduced by an additional 20% in size. **c.** The sensitivity and specificity of *ScoreRGM* estimated from the same simulation as in b. **d.** Likely mechanism of action for monotherapies used for treatments. **e.** Timeline of the experiment. Tumors were initiated in *KT*;*H11^LSL-Cas9^* mice with the barcoded Lenti-sg*TS*^Pool^/Cre. Three weeks of treatment was initiated after 15 weeks of tumor growth. **f-h**. *Ĝ* calculated from the inverse variance weighted average of *ScoreRTN* and *ScoreRGM* for the pharmacogenomic mapping experiment (**f**), negative control experiment in *KT* mice (**g**) and palbociclib repeat experiment (**h**). Stars represent significant cases. **i.** Comparison of our identified GSTRs with those from the Genomics of Drug Sensitivity in Cancer (GDSC) database. Stars represent significant cases. **j.** Kaplan-Meier curve (with 95% confidence interval in shading) of time-to-next-treatment (months) for patients with or without *KEAP1* mutations with metastatic oncogenic *KRAS*-driven lung adenocarcinoma to platinum-containing chemotherapy. The number of patients in each group is shown. *P*-values were calculated from the Mantel-Haenszel test. **k.** Responses of patients with metastatic oncogenic *KRAS*-driven lung adenocarcinoma to platinum-containing chemotherapy are consistent with *KEAP1* inactivation leading to resistance. *KEAP1* mutations are significantly correlated with a higher hazard ratio for time-to-next-treatment.

When quantifying tumor suppressor gene effects using Tuba-seq, each mouse represents an internally controlled experiment in which tumor size metrics can be normalized to tumors with inert sgRNAs within the same animal (**Fig. 1e, Supplementary Fig. 1**)^15^. In contrast, comparing tumor size distributions among groups of mice, such as between untreated and drug-treated groups, requires methods that overcome the technical and biological differences between mice. To address these challenges, we rigorously modeled drug responses and genotype-specific responses under the assumption that while tumors of all sizes respond equally to each treatment, the treatment effect can vary by genotype. Specifically, we estimated the drug effect on control tumors (those with inert sgRNAs) and then applied this effect to all tumors to calculate an expected distribution of tumor sizes after treatment (**Fig. 2a** and **Methods**). Genotype-specific therapeutic responses (GSTRs) were quantified by comparing the observed distribution of tumor sizes for tumors of a certain genotype after treatment with the expected distribution derived from the untreated mice. We developed two statistics to characterize GSTRs, the first is based on the relative numbers of tumors above a certain size after treatment (*ScoreRTN* – Relative Tumor Number) and the second compares the geometric mean of tumors from the full distribution of tumor sizes (*ScoreRGM* – Relative Geometric Mean) (**Methods**). By assessing the performance of the two statistics, we showed that both statistics are unbiased and have substantial and similar power although one statistic may outperform the other if the genotype-specific response is not uniform across tumor sizes (**Methods, Fig. 2b-c** and **Supplementary Fig. 3, 4** and **5**).

We applied Tuba-seq and our statistical metrics to assess the genotype-specific therapeutic responses of 11 genotypes of lung tumors to a panel of eight single and combination therapies (**Fig. 1a** and **2d**). These therapies were chosen to perturb diverse signaling pathways and assess the genotype-dependency of chemotherapy responses. *KT;H11^LSL-Cas9^* mice with Lenti-sg*TS^Pool^*/Cre-initiated lung tumors were treated for three weeks with one of the eight therapies followed by Tuba-seq analysis (**Fig. 1a**). The total cancer cell numbers estimated by Tuba-seq were highly correlated with total tumor-bearing lung weights, which varied substantially among mice even within the same groups (**Supplementary Fig. 6a-c)**. Despite such mouse-to-mouse variation, analysis of the overall tumor burden and the number of tumors with inert sgRNAs identified significant overall effects of five treatments (**Supplementary Fig. 6**).

We compared the tumor size profiles of treated mice with those of untreated mice and calculated the *ScoreRTN* and *ScoreRGM* (**Supplementary Fig. 7**). For both statistics, we estimated the magnitudes of GSTR and the associated *P*-values. Across all genotypes and treatments, the two statistics were well-correlated in magnitude as expected under the model of proportional tumor responses (**Supplementary Fig. 7b;** *r* = 0.86, *P*=10^-46^). Among the 88 assessed genotype-treatment pairs, 20 and 17 significant GSTRs (*P* < 0.05) were identified by *ScoreRTN* and *ScoreRGM*, respectively. Of these, 19 genotype-treatment interactions were significant by one statistic (*P* <0.05) and at least marginally significant (*P* < 0.1) by the other (**Supplementary Fig. 7a, b; Table S1)**. We derived a composite measure of GSTR (*Ĝ*) with the magnitude estimated from the inverse variance weighted average of the two statistics (**Methods**, **Fig. 2f**). Analysis of genotype-specific effects across treatments highlighted similarities among tumor suppressors, including those of Lkb1 and Setd2 known to have redundant tumor suppressive effects^5^, while combination treatments clustered with their corresponding single therapies (**Supplementary Fig. 7c, d**). Power analysis showed that our findings were robust to the choice of inert sgRNAs, cancer cell number cutoff, and inaccurate estimation of drug effect (**Supplementary Fig. 8, 9**).

Only one of the detected GSTRs was known in advance – the resistance of Rb1-deficient tumors to the CDK4/6 inhibitor, palbociclib. This resistance is consistent with the biochemical features of this pathway and clinical findings in breast cancer and hepatocellular carcinoma ^16–18^ (**Supplementary Fig. 11f**). Our ability to rediscover this interaction serves as a positive control of our method and is consistent with the expectation that some pharmacogenomic interactions transcend cancer types.

To further test the performance of our experimental and statistical procedures, we performed two additional experiments. First, as a negative control for GSTR identification, we treated Cas9-negative *KT* mice with a combination of chemotherapy and Mek-inhibition (**Supplementary Fig. 10a**). This treatment led to a dramatic reduction in tumor sizes compared to untreated *KT* mice (**Supplementary Fig. 10b**). Only one false positive GSTR was identified (*ScoreRTN*, P = 0.03; *ScoreRGM*, P = 0.07) with a very weak magnitude of the effect (*Ĝ* = 0.093, while the minimum magnitude of significant GSTR in the main experiment was 0.108; **Fig. 2g**). Combined with the fact that none of the individual inert sgRNAs (sg*Neo1*, sg*Neo2*, sg*Neo3*, and sg*NT*) had significant effects by either metric for any of the treatments in our main pharmacogenomic mapping experiment, this experiment provided additional confidence in the veracity of the detected GSTRs (**Fig. 2f**, **g**).

Simulation suggests that these cohort sizes have substantial power (**Supplementary Fig. 4**); therefore, we next attempted to rediscover the genotype-palbociclib interactions. We initiated tumors in a similar, albeit somewhat smaller cohort of *KT;H11^LSL-Cas9^* mice with Lenti-sg*TS^Pool^*/Cre and repeated the palbociclib-treatment. Analyses of these mice again identified *Rb1* inactivation as a mediator of palbociclib resistance (**Fig. 2f, Supplementary Fig. 10**). *Smad4*-deficient tumors, which showed modest resistance in our initial experiment, showed nominal resistance in the repeat experiment (*Ĝ* = 0.167), although this interaction was not significant (*P*=0.17 and 0.20 for *ScoreRTN* and *ScoreRGM*, respectively).

While the positive and negative predictive values of cancer cell line studies are often questioned ^19^, the scale at which these *in vitro* studies can be performed has enabled the generation of drug response data across large panels of cell lines ^4, 20^. We compared our findings to the largest dataset of cell line-therapeutic responses (Genomics of Drug Sensitivity in Cancer; GDSC)^3^ in which all five of our monotherapies were assessed (paclitaxel, palbociclib, phenformin, everolimus/rapamycin, and trametinib)^3^. Among the genotype-treatment pairs assessed in both studies, nine had significant effects in our analysis, but only one of these genotype-treatment interactions was significant in GDSC (*Rb1*-palbociclib)(**Fig. 2i** and **Supplementary Fig 12**). Three of the significant genotype-specific resistances, which were only uncovered *in vivo,* involved *Smad4*-deficiency (**Fig. 2i**). Given the absence of microenvironment stimulation of the TGFβ/Smad4 pathway in culture, these findings suggest that targeting paracrine TGFβ signaling could reduce the efficacy of cancer therapies in some contexts ^21^.

Although most of the detected pharmacogenomic interactions we uncovered are novel, several lines of evidence derived from clinical and preclinical data are consistent with our observations. For instance, *Lkb1*-inactivation reduced sensitivity to mTOR inhibition in our data, which is supported by previous anecdotal data from the analysis of lung adenocarcinoma patient-derived primary cultures *in vitro* (**Supplementary Fig. 13**)^5^. The ultimate goal of our study is to find genotype-treatment responses that predict lung adenocarcinoma patient responses. Lung adenocarcinoma patients are often treated with first-line platinum-containing combination therapies. In our analysis, *Keap1*-inactivation specifically led to resistance to treatments that included carboplatin, while not promoting significant resistance to the other therapies (**Fig. 2f**). To investigate the clinical impact of tumor suppressor genotype on lung adenocarcinoma responses, we queried the tumor suppressor genotypes and therapeutic benefit of platinum-containing treatments (assessed as time-to-next-treatment) of 216 oncogenic *KRAS*-driven human lung adenocarcinoma patients treated at Memorial Sloan Kettering Cancer Center (**Methods**). When each gene was assessed individually, both *KEAP1* and *LKB1* mutations were associated with worse clinical outcomes (*P*=6×10^-6^, **Fig 2j** and *P* = 0.06, **Supplementary Fig. 14j**, respectively). However, the marginally significant effect of *LKB1* mutation appears to be driven by the co-occurrence of *KEAP1* mutations with *LKB1* mutations (**Supplementary Fig. 14)**. This is also well supported by our pharmacogenomic data in which *Lkb1*-inactivation did not confer resistance to platinum-containing treatments (**Fig. 2f**). We further quantified the hazard ratio of the mutational status of the 11 genes in a manner that takes into account the effect of other co-incident mutations. This analysis confirmed that mutation of *KEAP1* correlated with a shorter time-to-next-treatment, which is consistent with our Tuba-seq results as well as a previous study on the impact of *KEAP1/NRF2*-pathway alterations on platinum responses (**Fig. 2j, k**)^22, 23^. Our *in vivo* pharmacogenomic platform, in which the responses of tumors with defined genotypes can be quantified, establishes direct causal relationships between genotype and treatment responses and enables accurate interpretation of patient data.

We uncovered a surprisingly complex pharmacogenomic map of resistance and sensitivity of *KRAS*-driven lung adenocarcinoma. Across diverse tumor suppressor genotypes and treatments, genotype-treatment interactions were very common (∼20% in our study), suggesting that genetic differences among tumors may underlie much of the heterogeneity in treatment responses. Our study required only a small number of animals, thus highlighting the power of our platform in confirming existing, and prioritizing new, genotype-specific drug responses (**Supplementary Fig. 15**). The exploitation of this platform to quantify the effects of additional therapies (and combination therapies) across a greater diversity of cancer genotypes will provide a cause-and-effect pharmacogenomic framework from which novel biological hypotheses and precision treatment approaches will emerge.

## Acknowledgements

We thank the Laura Andrejka for technical support; A. Orantes for administrative support; member of the Stanford Animal care staff for excellent animal care; D. Feldser, J. Sage, J. Zhang, and members of the Winslow and Petrov laboratories for helpful comments. C.L. is the Connie and Bob Lurie Fellow of the Damon Runyon Cancer Research Foundation (DRG-2331). W-Y.L. was supported by an American Association of Cancer Research Postdoctoral fellowship (17-40-18-LIN). H.R. was supported by the Druckenmiller Center for Lung Cancer Research at MSK. H.C. was supported by a Tobacco-Related Disease Research Program Postdoctoral Fellowship (28FT-0019). C.D.M was supported by NIH K99-CA226506. M.Y. was supported by a Stanford Dean’s fellowship, an American Lung Association senior research training grant, and NIH F32-CA236311. I.P.W was supported by the NSF Graduate Research Fellowship Program and NIH F31-CA210627. This work was supported by NIH R01-CA207133 (to M.M.W and D.A.P.), NIH R01-CA231253 (to M.M.W and D.A.P), NIH R01-CA234349 (to M.M.W and D.A.P.) and Memorial Sloan Kettering Cancer Center Support Grant/Core Grant (P30 CA008748).

## AUTHOR CONTRIBUTIONS

C.L., W.L., M.M.W., D.A.P., and C.M.R. planned the project. C.L. performed bioinformatics and statistical analysis. W.L., H.C., M.M.W., Z.R., and M.Y. performed the mice experiment. H.R. collected the patient response dataset. H.R. and C.L. performed the analysis on human dataset. Z.N.R., I.P.W., and C.D.M. assisted in the mice experiments. C.L., M.M.W and D.A.P. wrote the paper.

## COMPETING INTERESTS

D.A.P., I.P.W, and M.M.W are cofounders of D2G Oncology Inc. D.A.P., I.P.W, Z.N.R, C.D.M and M.M.W hold equity in D2G Oncology Inc.

## MATERIALS AND METHODS

### Generation of lentiviral vectors

The design and barcoding of Lentiviral sgRNA/Cre vectors were previously described ^15^. Briefly, each vector contains a guide RNA (sgRNA) targeting a specific tumor suppressor gene, a highly diverse barcode region (BC) (total length 23 bp, the length of the diverse sequence is 15bp), and an sgID sequence of 8 bp that is specific to each sgRNA. The BC and sgID regions are adjacent to one another and can be amplified in a single PCR reaction. To eliminate sgID-BC-sgRNA uncoupling driven by lentiviral template switching during reverse transcription of the pseudo-diploid viral genome, we produced each barcoded Lenti-sgRNA/Cre vector separately. We transfected 293T cells with individual barcoded Lenti-sgRNA/Cre plasmids (*sgApc, sgAtm, sgArid1a, sgCdkn2a, sgKeap1, sgLkb, sgNeo1, sgNeo2, sgNeo3, sgNT1, sgp53, sgRb1, sgRbm10, sgSetd2,* and *sgSmad4*) and packaging plasmids (delta8.2 and VSV-G) using polyethylenimine-based transfection. The supernatant was collected at 48 and 72 hours after transfection. Each lentiviral vector was filtered and concentrated by ultracentrifugation (25,000g for 1.5 hours), resuspended in PBS, and frozen in aliquots at −80℃. To determine the titer of each vector, we transduced LSL-YFP cells (a gift from Dr. Alejandro Sweet-Cordero/UCSF), determined the percent YFP-positive cells by flow cytometry, and calculated the titer by comparing to a lentiviral preparation of a known titer. Viral vectors were thawed and pooled immediately prior to delivery to mice.

### Mice, tumor initiation, and drug treatment

*Kras^LSL-G12D^ (K), R26^LSL-Tomato^ (T), H11^LSL-Cas9^* mice (hereafter *KT;H11^LSL-Cas9^* mice) have been previously described ^24–26^. Lung tumors were initiated using a pool of barcoded Lentiviral-sgRNA/Cre vectors delivered by intratracheal transduction of mice as previously described ^15^. 1.1 x 10^5^ and 2.2 x 10^4^ infectious particles in 60 μl of PBS were administered to each *KT* and *KT;H11^LSL-Cas9^* mouse, respectively. Tumor burden was assessed by fluorescence microscopy, lung weight measurement, and histology. Drug treatments were started 15 weeks after tumor initiation.

In the main pharmacogenomic mapping experiment, mice were assigned to eight treatment arms or 8 mice were left untreated for 3 weeks. There were five arms of monotherapies. 6 mice were treated with 100 mg/kg palbociclib daily by oral gavage; 5 mice were treated with 10 mg/kg everolimus daily by oral gavage, 6 mice were treated with 100 mg/kg phenformin daily by oral gavage, 5 mice were treated with 20 mg/kg paclitaxel every other day by intraperitoneal injection, and 7 mice were treated with 0.3 mg/kg trametinib daily by oral gavage. For mice treated with drug combinations, the dosing was the same as when each drug was used for monotherapies. 8 mice were treated with the combination of 20 mg/kg paclitaxel every other day by intraperitoneal injection and 0.3 mg/kg trametinib daily by oral gavage; 7 mice were treated with 50 mg/kg carboplatin every 5 days by intraperitoneal injection in combination with 20 mg/kg paclitaxel every other day by intraperitoneal injection; and 6 mice were treated by a combination of three drugs: 50 mg/kg carboplatin every 5 days by intraperitoneal injection, 20 mg/kg paclitaxel every other day by intraperitoneal injection, 0.3 mg/kg trametinib daily by oral gavage.

For the palbociclib repeat experiment, 5 mice were left untreated and 5 mice were treated with 100 mg/kg palbociclib daily by oral gavage. For the negative control experiment, 4 *KT* mice were left untreated, and 5 *KT* mice were treated with a combination of carboplatin, paclitaxel, and trametinib with the same dosing as the main experiment, except that treatment was continued for 6 weeks. All mice in all experiments were transduced with the same pool of viral vectors. All experiments were performed in accordance with Stanford University Institutional Animal Care and Use Committee guidelines.

### Isolation of genomic DNA from mouse lungs, preparation of sgID-BC libraries, and quantification of tumor size

Genomic DNA was isolated from bulk tumor-bearing lung tissue from each mouse as previously described ^15^. Three benchmark “spike-in” cell lines were added to each sample prior to lysis and DNA-extraction. Spike-in cell lines harbor integrated barcoded Lenti-Cre vectors with the sgID “TTCTGCCT”, which is distinct from all sgIDs in the Lenti-sgRNA/Cre pool. A pool of three Spike-in cell lines was added to each bulk lung sample at a known cell number (∼5×10^5^ cells per cell line), thus enabling the calculation of the absolute cancer cell number within each tumor ^14^. To determine the absolute number of cancer cells in each tumor, we multiplied the number of reads of each tumor sgID-BC by the mean number of reads of the three spike-in cell lines divided by the expected number of cells for each spike-in (5×10^5^ cells).

Libraries were prepared by PCR amplification of the sgID-BC region from 32 µg of genomic DNA per mouse. To enable the identification and subsequent computational elimination of possible index-hopped reads in high-throughput sequencing, the sgID-BC region of the integrated Lenti-sg*RNA-BC/Cre* vectors was PCR amplified from each sample with unique dual indexed primer pairs ^27^. Because Illumina short-read sequencing platforms require high library sequence diversity for accurate base calls, we previously added 50% PhiX phage DNA to each amplicon library before sequencing as recommended by Illumina. To further improve sequencing quality and minimize the use of PhiX DNA, we improved our previous method by adding 6 to 9 random nucleotides (Ns) to the flanking ends of both index primers before the sequence-specific primers. These random bases increase sequence diversity through their high nucleotide diversity and by desynchronizing the barcode regions of the amplicon library (via differing 6-9 nucleotide offsets). This reduced the needed PhiX genomic DNA from 50% to 10% and greatly improved Q30 scores.

Forward primers were of the form:

5’-AATGATACGGCGACCACCGAGATCTACAC**AGCGCTAG**

*ACACTCTTTCCCTACACGACGCTCTTCCGATCT* [N]_6-9_<u>GCGCACGTCTGCCGCGCTG</u> −3’

Reverse primers were of the form:

5’-CAAGCAGAAGACGGCATACGAGAT**CGTGAT**

*GTGACTGGACTTCAGACGTGTGCTCTTCCGATCT* [N]_6-9_<u>CAGGTTCTTGCGAACCTCAT</u> −3’

The underlined sequences denote regions that bind the template; the bolded sequences are the dual indexes which are unique to each mouse library pooled on the same sequencing lane; the italicized regions are the Illumina^®^ TruSeq Universal adapter sequence, and the sequences 5’ of the Indexes are the P5 and P7 adaptors.

We used a single-step PCR amplification of sgID-BC regions. We performed eight 100 µl PCR reactions per sample (4 µg DNA per reaction) using Q5 High-Fidelity 2x Master Mix (New England Biolabs)^28^ to maximize library sequencing quality. Pooled PCR products were isolated by gel electrophoresis and gel extracted using the Qiagen^®^ MinElute Gel Extraction kit. The concentration of purified PCR products from each sample was determined by Bioanalyzer (Agilent Technologies) and pooled for sequencing. Pooled libraries were cleaned up and size-selected using AMPure XP beads (Beckman Coulter). Libraries were sequenced on an Illumina^®^ HiSeq 2500 to generate paired-end 150 bp reads. Unlike patterned flow cells (e.g., HiSeq 4000), nonpatterned flow cells used by HiSeq 2500 exhibit extremely low index hopping rates^29^. Paired-end sequencing was used to ensure the fidelity of our barcode calls.

### Processing reads to identify the sgID and barcode

By systematically improving DNA library preparation and sequencing, we dramatically improved the quality of our barcoded tumor calls. In doing so, our previous DADA2-based denoising approach became obsolete, and a new, more stringent computational analysis pipeline that completely eliminated spurious tumors from PCR and sequencing errors was developed.

Read sequences are expected to contain an 8-nucleotide sgID region followed by a 23-nucleotide BC (AANNNNNTTNNNNNAANNNNN). We required both the forward and reverse sequencing reads to match perfectly within the BC region. This stringent requirement minimizes sequencing errors that will generate sgID-BC reads that do not represent actual clonal tumors within the samples. The FASTQ files of each technical replicate were combined for all analyses except when used for evaluating the reproducibility and quantifying error.

FASTQ files were processed to identify the sgID and BC counts for each tumor. The sgID region identified the targeted tumor suppressor gene, which we identified from the forward read alone, again allowing no mismatches or indels relative to the expected sgID sequences. Note that all sgID sequences differ by at least 2 nucleotides from each other, making this step robust to sequencing errors. The number of reads with each unique sgID-BC in each sample were added up to arrive at the size of each putative tumor (in the units of the number of reads).

### Identification of unique tumors from random barcodes and removal of “spurious tumor” generated by read errors

PCR and sequencing errors within the random barcode regions may be misinterpreted as unique tumors. These spurious tumors bias the analysis of tumor size distribution, reduce signals of tumor suppression, and potentially confound our ability to detect GSTR using Tuba-seq. Therefore, we used stringent criteria to reduce and even eliminate the effects of PCR and sequencing errors on tumor calls.

The random barcode in the sgID-BC region is 15 nucleotides long. Thus, there is a theoretical potential diversity of ∼4^15^ > 10^9^ barcodes within each lentiviral vector. While the actual diversity in each Lenti-sgRNA/Cre vector is dictated by the number of colonies generated during the plasmid barcoding step, this theoretical diversity guarantees that any two genuine tumors will be very unlikely to have barcodes within a certain Hamming distance from one another. Only approximately one pair of true tumors is expected to have barcodes that are only two nucleotides different from each other when making all pair-wise comparison in a set of 1000 tumors. Thus, we expect that most reads within two nucleotides from each other within the same mouse library are most likely due to sequencing/PCR errors. Therefore, we designed a pipeline in which any “tumor” with a barcode that was within a Hamming distance of two from a larger tumor in the same sample was excluded from subsequent analysis. As anticipated, excluded tumors were greatly enriched at the lowest end of the tumor size distribution, and were most often orders of magnitude smaller than the larger tumor with a similar barcode (Mann-Whitney U test of tumor sizes in the comparison between the removed and the remaining tumors, P<10^-300^). Thus, we elected to remove such putatively spurious tumors from subsequent analysis. Previously, we combined the smaller “spurious” tumors into the larger tumor with the similar barcode, as PCR and sequencing errors were more common and constituted a non-negligible fraction of the sequenced library. However, with our improved approach, these spurious tumors constituted only ∼ 0.3% of all reads and are necessarily more likely to be distinct tumors (as spurious tumors are now rarer), so we did not merge spurious tumors into their larger partners in this analysis. Because of the very small quantity of spurious tumors in our new approach, the decision to merge (or not merge) tumors have negligible effects on tumor size estimates.

To determine whether our stringent filtering removed most spurious tumors, we first estimated a False Positive Rate using our spike-in cell lines. We assessed whether any spurious tumors were generated from our spike-in controls for which we know the exact sequence of the sgID-BC. In these spike-ins, any sgID-BC read (i.e., with the spike-in sgID) containing a BC sequence that differs from the three known correct BCs is presumably generated by PCR or sequencing errors. Hence, these anomalous spike-in barcodes can be used to estimate the occurrence of spurious tumors. Before applying our filtering pipeline, we detected many such spurious barcode reads – 843 spurious spike-in “tumors” in a typical sequencing lane (**Fig. 1c**). Our new stringent filtering removed all of them.

We next identified true positive tumors by sequencing the sgID-BC region in each plasmid pool used to generate the lentiviral vectors in the Lenti-sgTS^Pool^/Cre pool. Given the tremendous theoretical diversity of our random barcodes, barcodes detected in *both* the plasmid pool and the mouse lungs are very likely to be genuine. We used these data to assess the presence of spurious tumors in the mouse lung as these should almost never be detected in the plasmid pool. However, a false positive rate cannot be directly measured from this plasmid sequencing data because barcode diversity in the plasmid pool is too high to be exhaustively sequenced to cover every potential barcode, i.e., not all barcodes present in the plasmid pool were captured in the barcodes sequenced from the plasmids. Thus, some genuine tumors in the mice will have barcodes not captured in the plasmid sequencing. Nevertheless, in a well-calibrated analysis pipeline, the rate at which genuine tumors are detected should not vary with tumor size. We determined whether the smaller tumors – which are more likely to be polluted by spurious tumors – exhibit a lower probability of detection in the plasmid pool. Before the application of our filtering procedure, 92.1% of tumors larger than 1000 cells, but only 87.9% of tumors smaller than 1000 cells were detected in the plasmid pool (Chi-squared test, p<10^-300^). This decline in the true positive tumor fraction with tumor size suggests that some spurious small tumors exist in our data prior to filtering. However, after our filtering procedure, 92.7% of tumors larger than 1000 cells and 93.1% of tumors smaller than 1000 cells were detected in the plasmid pool (**Fig. 1d** and **Supplementary Fig. 1b**). This consistency of the true tumor rate across tumor size suggests that filtering effectively eliminated spurious tumors. Along with our analysis of spurious spike-ins, both results suggest that our filtering procedure effectively eliminates spurious tumors.

### Summary statistics for characterizing tumor growth

Because GSTR can manifest as a mean effect or a change in the shape of tumor size distributions, we used a variety of summary statistics to characterize tumor growth, including percentiles, Log-normal (LN) mean, geometric mean, and relative tumor burden. Percentiles and LN mean were calculated as previously described ^15^. Briefly, percentiles are nonparametric summaries for the distribution. 95^th^ percentile for *sgTS* tumors, for instance, is calculated as the size above which we find the top 5% of the largest *sgTS* tumors and then dividing it by the corresponding number for the tumors with inert sgRNAs (Inert tumors). The resulting number is dimensionless. The LN mean is the maximum likelihood estimator of the mean number of neoplastic cells given a log-normal distribution of tumor sizes. The LN mean of sgTS tumors is relative to the LN mean of the Inert tumors. The geometric mean is defined as the product of all tumor sizes raised to the inverse of the number of tumors. For calculation, we compute the arithmetic mean of the logarithm-transformed values of tumor sizes and then use the exponentiation to return the computation to the original scale. The geometric mean is proportional to the average growth rate of tumors. Relative tumor burden is calculated as

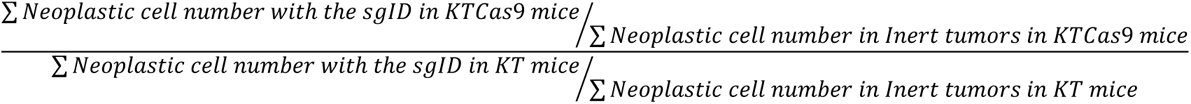

### Developing unbiased procedures for the detection of genotype-specific drug effects

Previous Tuba-seq analyses focused on comparing the sizes of tumors of different genotypes within individual mice ^14, 30^. Such analyses are largely robust to multiple sources of variation among mice such as (1) variation in the efficiency of viral delivery and the resulting differences in tumor number and total tumor burden across mice and (2) variation in the library sequencing depth (each sgID within a mouse is part of the same DNA library reaction). However, in the current analysis of genotype-specific drug response, we needed to compare tumor sizes between the two groups of mice – the untreated and the treated – rather than only comparing the relative behaviors of tumors of different genotypes within mice and then aggregating the signals across mice. Thankfully, our multiplexed pool of tumor genotypes still allows us to account for variations among mice, as these effects are still common to all imparted genotypes. First, however, we must generate a genotype-nonspecific null model of response. Note that because we used the same viral pool to initiate tumors in all treated and untreated mice, the initial relative representation of transduced epithelial cells containing each Lenti-sgRNA/Cre is constant and does not vary across mice. Nevertheless, our null model does not assume that the total numbers of initiated tumors among mice are invariant; only the *proportions* of initiated tumors with different sgRNAs (sgIDs) remain constant.

### Null model of tumor responses with no genotype-specificity

We assume that the therapy affects all tumors proportionally to their sizes (proportional size reduction) such that the size of each tumor changes from X to X_1_ = X × *S* after the drug treatment, where *S* is the proportion of remaining cancer cells. In other words, we assume that the therapy kills individual cancer cells with a probability 1-*S* that is independent of the tumor size. Under the null model (H_0_) of no genotype-specific drug responses, *S* is constant and does not depend on the genotype of the tumor. Under the alternative model H_1_, *S* varies depending on the genotype: *S*_sgID, j_ = *S*_Inert_×(1+*G*_j_), with *G_j_* representing the Genotype Specific Therapeutic Response (GSTR) of tumors generated by viruses with the specified sgID to the drug j. If *G*_j_ > 0, the inactivation of the tumor suppressor associated with that sgID confers relative resistance; if *G*_j_ < 0, the inactivation of the tumor suppressor associated with that sgID confers relative sensitivity.

### Selection of a size cutoff in untreated mice

To consider how tumor size distributions shift between the null and alternative models, we must first choose the range of the tumor size distribution to consider. In general, we want to consider tumors that are large enough to be consistently detected regardless of the sequencing depth and PCR efficiency in both treated and untreated mice, while using as many tumors as possible to maximize the statistical power. As shown below, the most extreme treatment reduced tumor sizes by ∼87% (*S*=0.13, **Supplementary Fig. 6d**). While the depth of sequencing varies across mice and treatments, we want to reliably identify tumors in each treated and untreated mouse. The most effective treatment group has the smallest tumors after treatment, for which the depth of sequencing was ∼10 cells/read. Thus, we chose to use the cutoff of *L* = 1000 cells in the untreated mice, as tumors will not shrink below 100 cells or 10 reads in all treated mice, allowing reliable detection and accurate size estimates of tumors in each mouse. **Supplementary Fig. 8** shows that our results are robust to shifting the cutoff to 500 or 1500 cells.

### Calculation of proportional size-reduction as the drug effect

We first find the value of the tumor reduction factor *S* that leads to the best match between the distributions of Inert tumors between the treated and untreated group under our model of proportional tumor reduction. We take the following steps to calculate *S*. For each possible value of *S* between 0 (tumors are completely eliminated) and 2 (tumors doubled in size after treatment), we reduce the sizes of each tumor in the untreated group by *S* and calculate the number of such “shrunk” tumors whose sizes remain above or equal to 1000 cells. We then find the value of *S* such that the median number of such shrunk tumors across all the untreated mice is closest to the median of the number of observed tumors with the size above or equal to 1000 cells across all the mice in the treated group. Specifically, we use the binary search algorithm to determine the *S* that minimizes the difference between the median numbers of tumors of the treated and untreated groups. We choose to find the *S* that matches the median number of tumors, rather than the mean across the two groups, as the median is not affected by the outlier mice with very low or very high numbers of tumors. Since we have prior knowledge that drugs will not increase overall tumor size, an estimated *S* larger than 1 is probably due to mouse-to-mouse variations. Thus we set *S* to 1 when it was estimated to be larger than 1. In **Supplementary Fig. 9,** we showed that our estimation is robust to the inaccurate estimation of *S*, and the power will only be reduced slightly when *S* was not accurately estimated.

### Approach 1: Relative tumor number (*ScoreRTN*)

Our first approach defines response as the number of tumors that exceed a minimum size threshold. The intuition is that, given a known tumor reduction factor *S* of the drug, the null hypothesis for each genotype is that the number of tumors above the cutoff *L* in the untreated mice should match the number above the new cutoff *L*×S in the treated mice. If a GSTR exists (the alternative model), then the tumors with a specific sgID (*i.e.,* tumors with a particular tumor suppressor inactivated) are more resistant to the drug than the Inert tumors and more of such tumors should remain above the adjusted cutoff of *L*×*S* than expected, while if they are more sensitive, then fewer of such tumors should remain above the adjusted cutoff of *L*×*S* – rejecting the null hypothesis in either case.

To test this null hypothesis, we first calculate the ratio of the number of tumors above the cutoff *L* in the untreated mice of a particular sgID to that of the Inert tumors (*RTN*_i,j,L_),

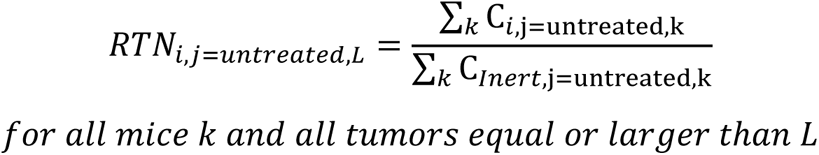

where *C*_*i*,*j*,*k*_ is the total number of tumors observed in mouse *k* in treatment group *j* (*j* = untreated here) carrying sgID *i* above the cutoff *L*. We then calculate the similar ratio for the treated mice with a modified cutoff *L*×*S*,

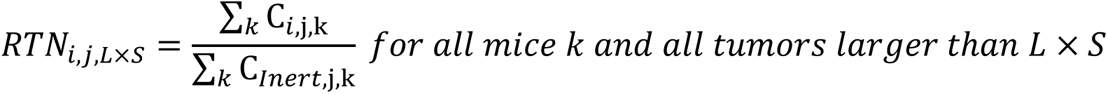

The null hypothesis can then be expressed as the expectation that

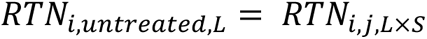

or alternatively that:

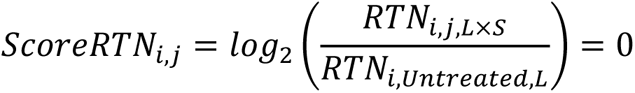

Under the alternative hypothesis where *ScoreRTN*_*i*,*j*_ ≠ 0, a positive sign of *ScoreRTN*_*i*,*j*_ suggests that the tumors with a particular sgID are more resistant than the Inert tumors, while a negative sign suggests the tumors are more sensitive than Inert tumors.

Although directly comparing the size or number of tumors above a constant cutoff (e.g.,1000 cells), for both the untreated and treated groups may seem intuitive and simpler, such a comparison generates complex expectations of tumor number that depend both on the distributions of tumor sizes prior to treatment and the magnitude of the drug effect (**Supplementary Fig. 3**).

### Approach 2: Relative geometric mean (*ScoreRGM*)

The second metric, Score*RGM*, compares the geometric mean of tumors carrying sgID *i* relative to the Inert tumors in the untreated and treated groups. The intuition is that if we analyze a comparable number of tumors in the untreated and treated mice when there is no *GSTR*, the relative growth advantage of tumors carrying a specific sgID (sgID *i*) relative to Inert tumors, represented by the relative geometric mean, will remain constant. Under the alternative model, if the tumors with a specific sgID (sgID *i*) are more resistant to the drug than the Inert tumors, then the relative geometric mean for sgID *i* will be larger in the treated group, while if they are more sensitive, then the relative geometric mean for sgID *i* will be smaller. While *RTN* does not use the numeric value of tumor size other than comparing it with the cutoff, (*i.e.,* a tumor with size 1001 cells and a tumor with size 10^7^ cells are both counted as a single tumor above the cutoff of 1000 cells), *RGM* incorporates such tumor size profile information. Hence, *RGM* and *RTN* are not entirely redundant as they incorporate different information about *GSTR*. Based on power analysis, *ScoreRTN* is a more sensitive metric in detecting *GSTRs* (**Fig 2b, c,** and **Supplementary Fig. 4**), particularly when only smaller tumors show *GSTR* (**Supplementary Fig. 5a**). However, when only larger tumors show GSTR, the *ScoreRGM* is more likely to capture it, as large tumors are unlikely to fall below the given size threshold. For instance, in an extreme case, when only large tumors with over 4000 cells show resistance, *ScoreRTN* will fail to capture the signals that *ScoreRGM* identifies with reasonable power (**Supplementary Fig. 5b**).

We denote the total tumor count (*T*) with a certain sgRNA (*i*) in an individual mouse (*k*) in the treated group (*j*) as *T*_i,j,k_. Here, we do not limit tumors to those above 1000 cells but rather count any tumor with greater than or equal to 2 reads (after the stringent filtering described above) as a tumor. For an untreated mouse, the proportion of initiated tumors of each sgID can be approximated by *R*_i_, the ratio of *T_i,untreated,k_* to *T_Inert,untreated,k_*:

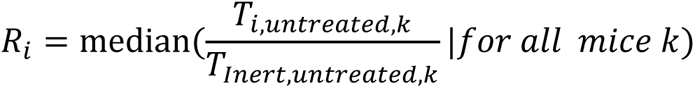

We then take the top *N* tumors with sgRNA *i* from mouse *k* treated by drug *j* as:

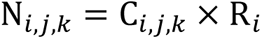

where *C*_*i*,*j*,*k*_ is the total number of Inert tumors observed in each mouse above the cutoff *L* × *S* (*S*=1 for the untreated group), and then we calculate the geometric mean for all tumors containing the sgID and Inert tumors across all mice in the group.

The score for the relative geometric mean is calculated as:

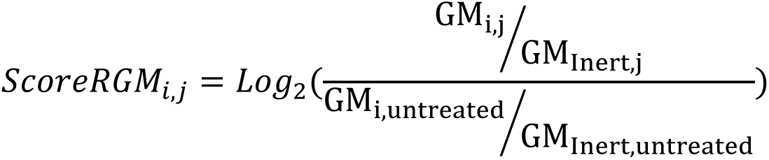

where GM_i,j_ is the geometric mean for tumors containing sgID *i* in treatment group *j* in the selected *N* tumors. Under the null hypothesis, *ScoreRGM*_*i*,*j*_ = 0. Under the alternative hypothesis where *ScoreRGM*_*i*,*j*_ ≠ 0, a positive sign of *ScoreRGM*_*i*,*j*_ suggests that the tumors with a particular sgID are more resistant than the Inert tumors, while a negative sign of the score suggests that these tumors are more sensitive than the Inert tumors.

### Evaluating whether *ScoreRTN* and *ScoreRGM* are biased

To test whether either statistic is biased, we calculate the *ScoreRTN* and *ScoreRGM* using untreated mice to subjected to a simulated treatment with no GSTR (specifically, the 5 untreated mice from the palbociclib repeat experiment). This represents an ideal scenario where we know exactly how each tumor responded to the treatment (because we generate the responses through simulations), and we are able to measure the exact tumors with and without treatment. Specifically, as shown in **Supplementary Fig. 3a**, we reduce the tumor sizes of all tumors in the five untreated mice by 50% as the “treated” tumors (**Supplementary Fig. 3b**). For a biased method illustrated in **Supplementary Fig. 3c, e, f,** which fails to consider the size reduction of tumors due to drug treatment by using the constant 1000 cell cutoff in both untreated and treated mice, the relative tumor number and relative geometric mean does not remain constant between treated and untreated mice (**Supplementary Fig. 3e, f**). This is because a constant cell number cutoff effectively compares different proportions of the distributions of the inert and TS-inactivated tumors in the treated and untreated mice.

On the other hand, using the adaptive cutoff method introduced above, no false signals of *ScoreRTN* and *ScoreRGM* were observed (**Supplementary Fig. 3d, g, h**), because we are comparing the matched portions of the distributions for the Inert tumors and tumors of each genotype (tumor with each sgID) between untreated and treated mice. Therefore, these two statistics appear unbiased.

### Bootstrapping the tumors

When generating null distributions of scores and calculating the p-values, we performed bootstrap resampling on tumors. During bootstrapping, we consider tumor size variations both across mice and within a mouse using a nested resampling approach: first, we bootstrapped mice in the untreated and/or the treated group to generate pseudogroups of mice, and then within each mouse, we bootstrapped all observed tumors carrying each sgID.

### Generate a null distribution of *ScoreRGM* and *ScoreRTN*

To generate a null distribution of *ScoreRTN* and *ScoreRGM* in the absence of GSTR, we sampled with replacement the same number of mice as in the treated group from the 8 untreated mice and applied estimated drug effects *S* on the tumors from the “treated” groups. For each bootstrap run, we re-estimated *S* and calculated the values of *ScoreRTN* and *ScoreRGM*.

### Generate the observed distribution and calculation of the confidence interval for each score

To calculate the confidence interval for each score, we bootstrapped mice in the treated and untreated groups, respectively, and then re-calculated *S*, *ScoreRTN,* and *ScoreRGM*. The bootstrap process was performed 10,000 times, and the 95% confidence interval of each score was calculated as the 2.5%ile, and 97.5%ile of the bootstrapped results.

### Calculation of the *P*-value for *ScoreRTN* and *ScoreRGM*

To see how the distribution of observed *ScoreRTN* and *ScoreRGM* of sgID *i* deviates from the null distribution, we compared the distribution of the two observed scores to that calculated from (1) simulated data with no *GSTR* for all sgIDs and to (2) simulated data with no *GSTR* for sgID *i* to determine the *P*-value. For each comparison, we sample values from both distributions and calculate the *P*-value as the fraction of times when their differences are not in the same direction as the observed score, *i.e*., how often do we see equal or more extreme scores under the null distribution. To be conservative, the maximum of the two *P*-values calculated from the two comparisons were reported as the *P*-value. This bootstrap process is performed 10^8^ times to calculate the *P*-values.

### Power analysis for *ScoreRTN* and *ScoreRGM* in our study

To estimate the sensitivity (True Positive Rate) and specificity (1-False Positive Rate) of our study (**Fig. 2b, c**), we sampled with replacement eight and five mice (minimum number of treated mice in the pharmacogenetics mapping experiment) from the eight untreated mice, as the “untreated” and “treated” groups, respectively. We then reduce all tumor sizes to 50% as the drug effect (*S*=0.5) and apply an input *G* = −50%, −20%, 0% (no genotype-specific drug response), 10%, 20%, and 50% to each non-Inert sgIDs by additionally changing each tumor sizes by the corresponding proportions. For instance, *G*=20% means the overall drug effect on tumors carrying the sgRNA is (1-50%)×(1+20%)=60%. Sensitivity is calculated as the probability of detecting preassigned true genotype-specific interactions, and specificity is calculated as the proportion of sgIDs correctly identified as having no *GSTR* when *G* = 0. A total of 100 runs of simulations of the 11 tumor suppressor gene targeting sgIDs were performed for each preassigned *G*. We adjust the cutoff for *P*-values and calculated a series of sensitivity and specificity values to plot the receiver operating characteristic (ROC) curve.

### Power analysis for *ScoreRTN* and *ScoreRGM* for various sample sizes

To evaluate the power of our method using various sample sizes (**Supplementary Fig 4**), we sampled the same number of mice (5, 10, or 20 mice) with replacement from the 8 untreated mice as the untreated and treated group. Similar to the previous section of Power analysis using 8 and 5 mice respectively for untreated and treated mice, we apply *G* of various magnitudes and plotted the ROC curve when we use 5, 10, and 20 mice for each group in our experiment, respectively.

### Power analysis for *ScoreRTN* and *ScoreRGM* for tumor size-dependent GSTR

We evaluate the effectiveness of *ScoreRTN* and *ScoreRGM* in capturing GSTR when the input *G* is not a constant factor constant across tumors with various sizes (**Supplementary Fig. 5**). We apply a truncated effect of *G* where 1) only tumors smaller than 4000 cells showed genotype-specific sensitivity or 2) only tumors larger than 4000 cells showed genotype-specific resistance. The rest of the tumors are simulated to respond the same as Inert tumors to the drug. We calculated the ROC curves for the two statistics using 5 mice in both the untreated and treated groups.

Apart from *ScoreRTN* and *ScoreRGM,* which are based on the relative tumor number and relative geometric mean, other summary statistics, such as **r**elative **L**N **m**ean (*ScoreRLM*), can also be used to identify GSTR. The score for the relative LN mean is calculated as:

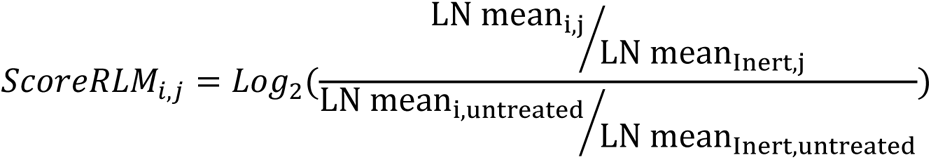

where LN mean_i,j_ is the LN mean for tumors containing sgID *i* in treatment group *j* in the selected *N* tumors. We compared and contrasted the performance of the three metrics when the GSTR is not a constant factor, and show that if we have a good reason to believe that only very large tumors show genotype-specific responses, *ScoreRLM* is the best metric among the three. Otherwise, the other two metrics, *ScoreRTN* and *ScoreRGM*, will outperform *ScoreRLM*.

### Evaluating whether inaccuracy of drug effect (*S*) estimation influences estimates of GSTR

We also evaluate the impact of inaccurately estimating drug effects on our estimates of GSTR (**Supplementary Fig. 9**). Instead of estimating the effect size as described in section “calculation of proportional size-reduction as drug effect”, we assign *S* to be a constant value, taking three discrete values 0.3, 0.5, and 0.7 when we know the simulated truth of drug effect is *S*=0.5. For each simulated scenario, we calculated the specificity and sensitivity and plotted the ROC curve for detecting *G*=+20% using 5 mice in both the untreated and treated groups (**Supplementary Fig 9**).

### Calculating ScoreGSTR (***Ĝ***) as the combined score

Although *ScoreRTN* and *ScoreRGM* may have an emphasis on different aspects of *GSTR* on tumor size distribution, it would be helpful to have a single combined score. We calculated a combined score of GSTR (*Ĝ*) by taking the inverse variance weighted average of *ScoreRTN* and *ScoreRGM*, then converting it to the linear scale (**Fig 2f, Supplementary Fig 4c).**

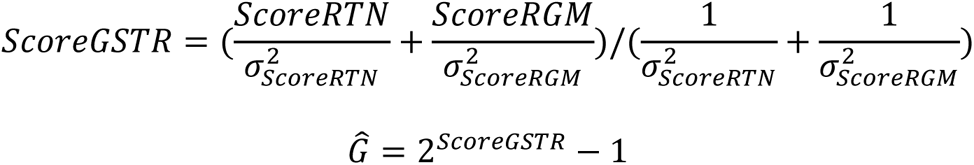

If 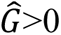, *GSTR* is resistant, and if 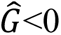, *GSTR* is sensitive.

To be very conservative, for the combined score to be called significant, we require at least one significant *P*-value (*P* < 0.05), and one marginally significant *P*-value (*P* < 0.1) for the two statistics *ScoreRTN* and *ScoreRGM*.

### Evaluate the consistency for choosing various cell cutoffs and control sgIDs

We use a cutoff of 1000 cells for most parts of the analysis, but we also wanted to evaluate whether our results are robust to using higher and lower cell number cutoffs. Therefore, we adjusted the cutoff to 500 cells and 1500 cells and re-identified the significant GSTR under each scenario (**Supplementary Fig. 8a**).

For most analyses, we aggregate tumors with sg*Neo1*, sg*Neo2*, sg*Neo3,* and sg*NT1* as a single Inert control sgRNA to determine the baseline. We also explored whether excluding any of the four sgIDs associated with Inert sgRNAs from the control group would yield similar results. For comparison, we calculated the Spearman correlation and linear correlation, identified significant cases of GSTR, and the overall direction of GSTR for each scenario compared with using all four sgIDs associated with Inert sgRNA.

### Comparing with human cell line response data

The drug sensitivity data from human cell lines were downloaded from the Genomics of Drug Sensitivity in Cancer (GDSC) database (www.cancerrxgene.org)3. GDSC used logic-based modeling to quantified how genetic alterations in 1001 human cancer cell lines are correlated with sensitivity to various drugs. There is a limited number of LUAD cell lines in the database. Therefore, we focused on comparing the results from Pan-cancer cell lines. All 5 monotherapies used in our study were assessed by GDSC. Except for *Keap1* and *Rbm10,* which are not reported for everolimus and paclitaxel, the GSTR of all other 51 gene-drug pairs were quantified by GDSC. The effect size and FDR-corrected *P*-values were used for comparison.

### Hierarchical clustering of *GSTR* for treatment and genes

To better visualize the similarity of the *GSTR* profiles, we performed hierarchical clustering with complete linkage on *Ĝ* to visualize the relationship across different genes and across different therapies (**Supplementary Fig 7c, d**). Therapies or genes that are similar to each other in genotype-specific responses are clustered together.

### Analysis of clinical data for resistance to chemotherapy

Patients with metastatic or recurrent lung adenocarcinoma harboring a KRAS mutation in codons 11, 12, or 61, as detected by MSK-IMPACT ^31^, were reviewed. Patients who received platinum chemotherapy (carboplatin or cisplatin) with pemetrexed +/-bevacizumab as first-line treatment were included (n = 216). Treatment efficacy was measured as time of first treatment with platinum doublet chemotherapy to start of next systemic therapy, or death if no subsequent therapy was received. Patients who continued on platinum doublet therapy at the last follow up were censored. Data collection was approved by the MSK institutional review board.

Kaplan-Meier estimator plots of time-to-next-treatment for patients with and without mutations at each of the 11 tumor suppressor genes of interest were generated. In addition, a multivariable Cox proportional hazards model analysis was performed integrating the mutational status of the 11 genes as individual input features to assess the independent effect of co-occurring mutations.

---

**Supplementary Figure 1.**
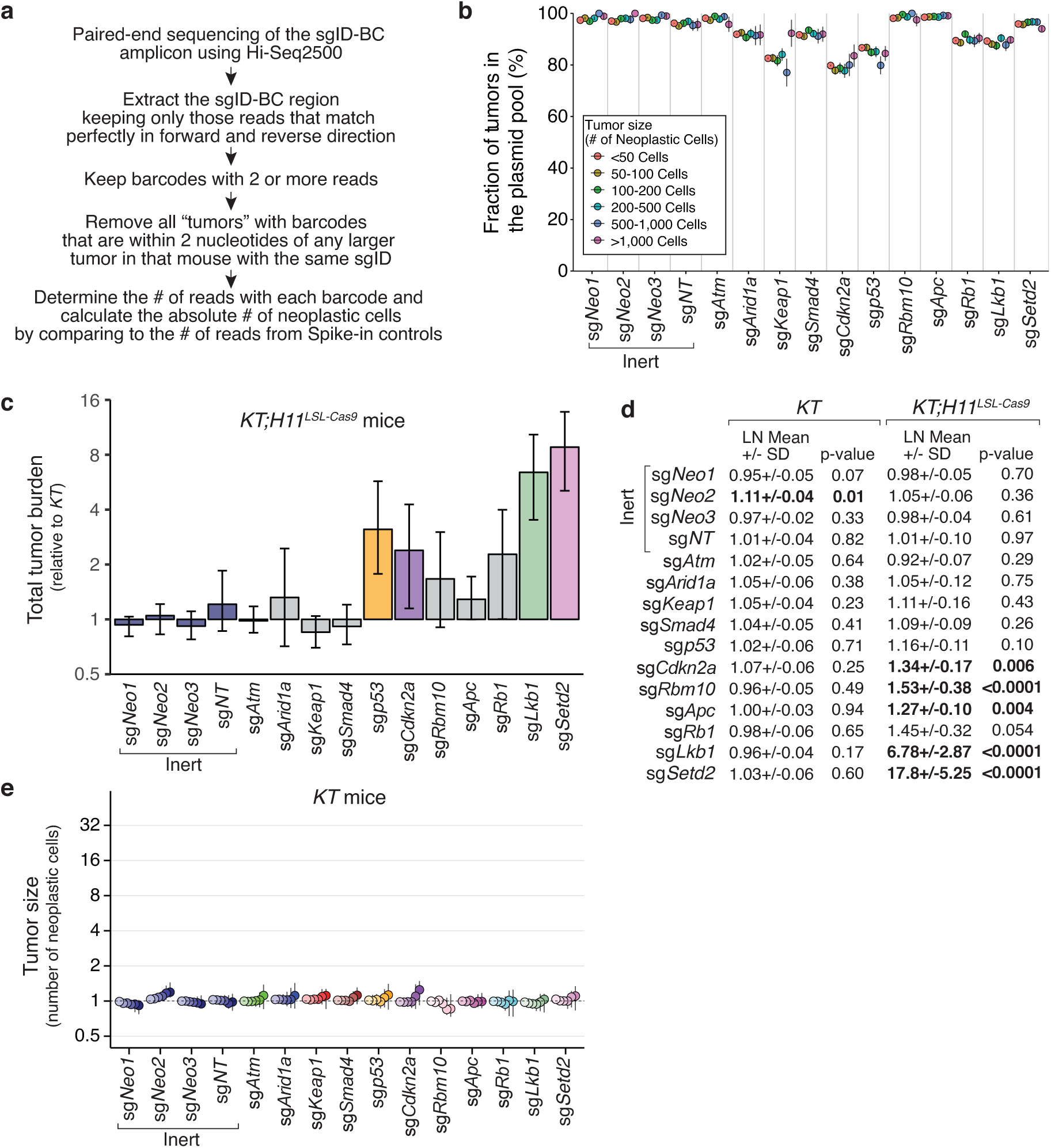
Optimization of Tuba-seq increases the resolution and precision of tumor analyses. **a.** Overview of our new Tuba-seq analysis pipeline for calling sgID-BC from sequencing data and determining the number of neoplastic cells in each tumor (tumor size). **b.** Fractions of tumor sgID-BC region recovered in the plasmid pool across multiple size ranges. By sequencing the sgID-BC region in the Lenti-sgRNA/Cre plasmids, we define a high confidence list of barcodes that are present in each sgID-BC region. If spurious tumor remains, the identified smaller tumors in *KT;Cas9* mice will less likely be uncovered in the plasmid pool compared with larger tumors. However, we find that equivalent proportions of tumors at all sizes are present in the plasmids pool. Note that not all barcodes found in tumors are found in the sequenced plasmids pool because the sequencing depth of the plasmid pool was insufficient to uncover all barcodes in the plasmid pools. **c.** Total tumor burden in *KT;Cas9* mice with Lenti-TS^Pool^/Cre-initiated tumors relative to expected tumor burden calculated from *KT* mice. Error bars show 95% percent confidence intervals. Note that targeting p53 enables the generation of rare very large tumors, hence the tumor-suppressive effect of p53 is easily identified by this metric (which aggregates all cancer cells), while the 95^th^ percentile of tumor size distribution and LN mean are much less dramatic. **d.** The relative LN mean for tumors with each sgRNA in *KT* and *KT;H11^LSL-Cas9^* mice 18 weeks after tumor initiation normalized to that of all Inert tumors. Bootstrapped p-values are shown. *P*-values < 0.05, and their corresponding means are in bold. **e.** The relative size of tumors initiated with each Lenti-sgRNA/Cre vector in *KT* mice 18 weeks after tumor initiation with Lenti-sgTS^Pool^/Cre. Relative size of tumors at the indicated percentiles within the distribution represent merged data from 4 mice. 95% confidence intervals are shown.

**Supplementary Figure 2.**
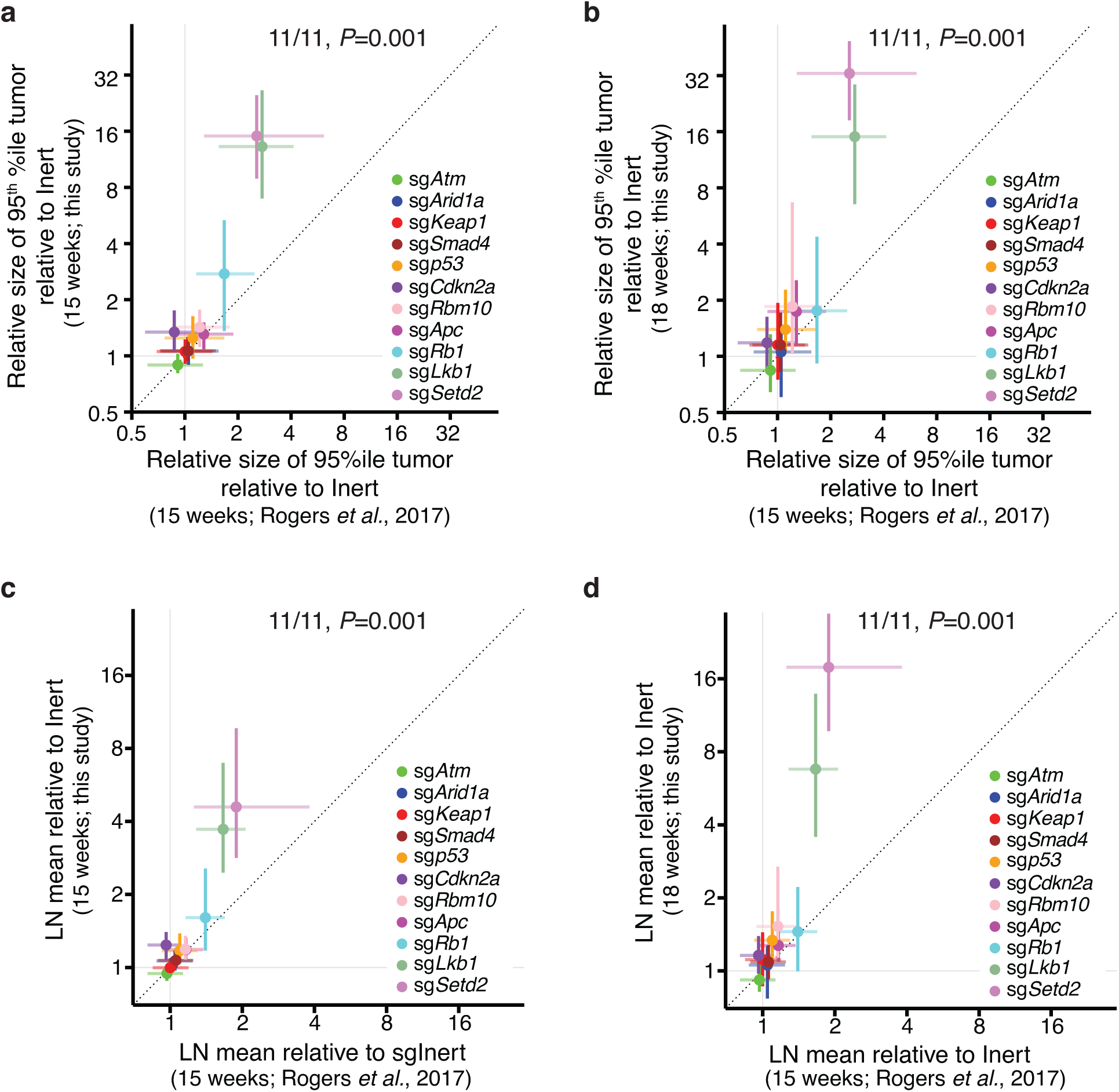
Optimization of Tuba-seq increases the resolution and precision of tumor analyses. **a,b.** Comparison of the relative 95^th^ percentile tumor sizes (size of 95^th^ percentile sgTS tumor/size of 95^th^ percentile Inert tumor) of each genotype between the previous data (Rogers *et al.*, 2017) and the current data (this manuscript). Error bars show the 95% confidence interval. Current data from tumors 15 weeks (**a**) and 18 weeks (**b**) after tumor initiation are shown. **c,d.** Comparison of the relative LN mean (LN mean of sgTS tumor/LN mean of Inert tumors) of the tumors of each genotype between the previous data (Rogers *et al.*, 2017) and the current data (this manuscript). Error bars show the 95% confidence interval. Current data from tumors 15 weeks (**c**) and 18 weeks (**d**) after tumor initiation is shown. In each panel, Error bars show the 95% confidence intervals. Dash lines represent equal detected magnitudes of tumor suppression across studies. 11/11 means 11 out of 11 the assayed genotypes showed a higher magnitude in current studies, and *P*-values are calculated from the sign-test of the difference in magnitudes for each metric between this study and the previous study.

**Supplementary Figure 3.**
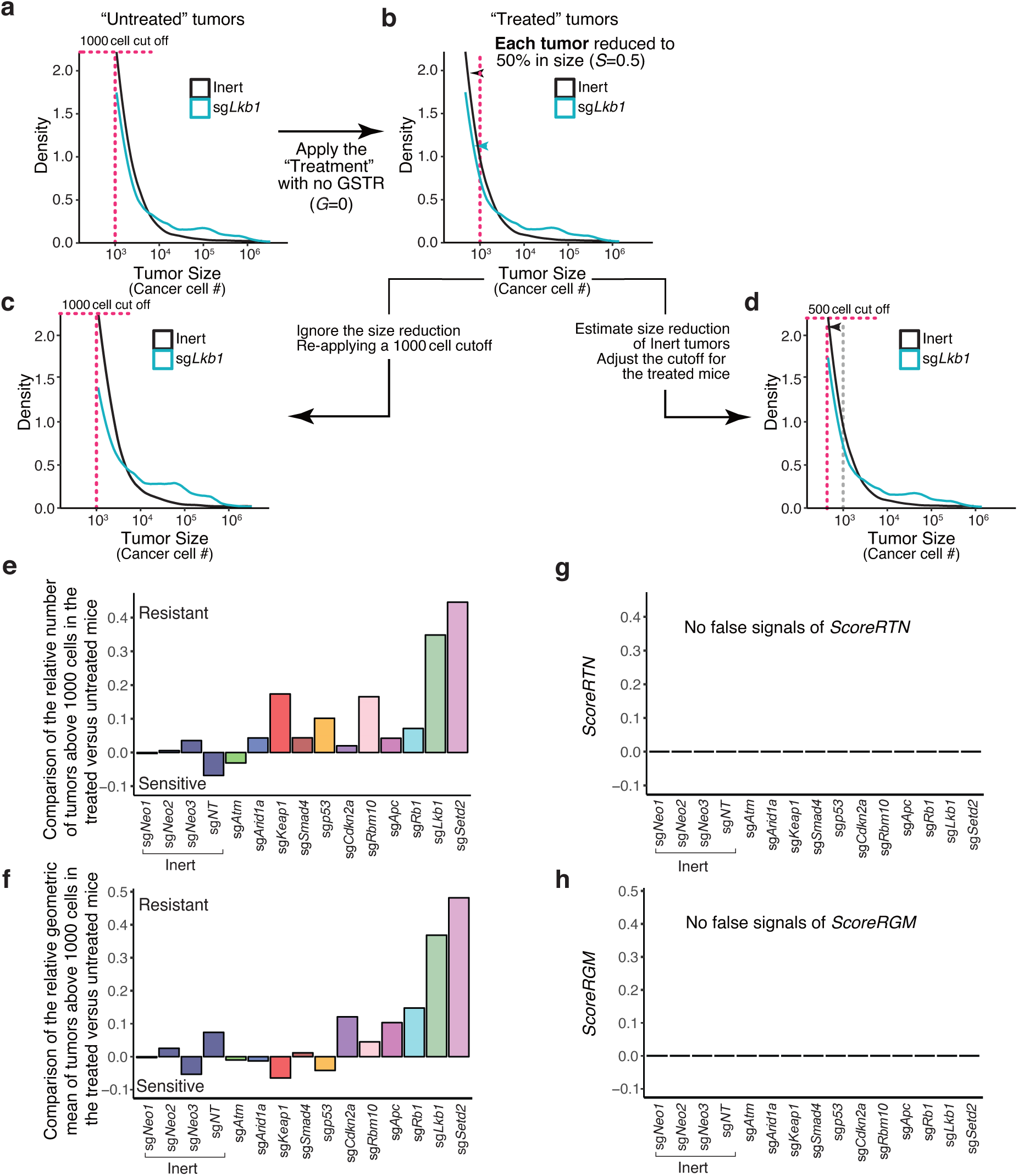
The intuitive approach of comparing tumors above a constant cutoff between the untreated and treated mice generates aberrant signals of Genotype-specific treatment responses (GSTRs) **a.** Tumor size distributions of “untreated” Inert and sg*Lkb1* tumors were pooled from 5 *KT;Cas9* mice 18 weeks after tumor initiation for all simulations. Tumors with more than 1,000 neoplastic cells are plotted. Inert tumors are tumors containing sg*Neo1*, sg*Neo2*, sg*Neo3*, or sg*NT*. Targeting *Lkb1* changes the overall shape of the tumor size distribution. **b.** Tumor size distributions of “treated” Inert and sg*Lkb1* tumors were generated by reducing the size of every tumor to 50% (*S*=0.5), assuming each cancer cell was killed by the drug with a 50% probability (*G*=0). **c.** After re-applying a 1000 cell cut off, the new tumor size distributions of Inert and sg*Lkb1* tumors are noticeably different from those before the size reduction in panel a. **d.** To account for tumor size reduction, we need to adjust the cutoff in the treated group by the estimated size reduction due to treatment. **e,f.** Comparing the relative tumor number (**e**) and relative geometric mean (**f**) (normalized to the corresponding Inert tumors) between the “untreated” and “treated” tumors with each sgID by taking the log_2_ ratio of the metric in the treated mice over that of the untreated mice for tumors with each sgID. The values are non-zero. Therefore, this intuitive approach is incorrect. **g,h.** After correcting for the overall tumor size reduction by the treatment, no false signals of ScoreRTN (**g**) and ScoreRGM (**h**) were generated when comparing the relative tumor number and relative geometric mean between the “untreated” and “treated” mice.

**Supplemental Figure 4.**
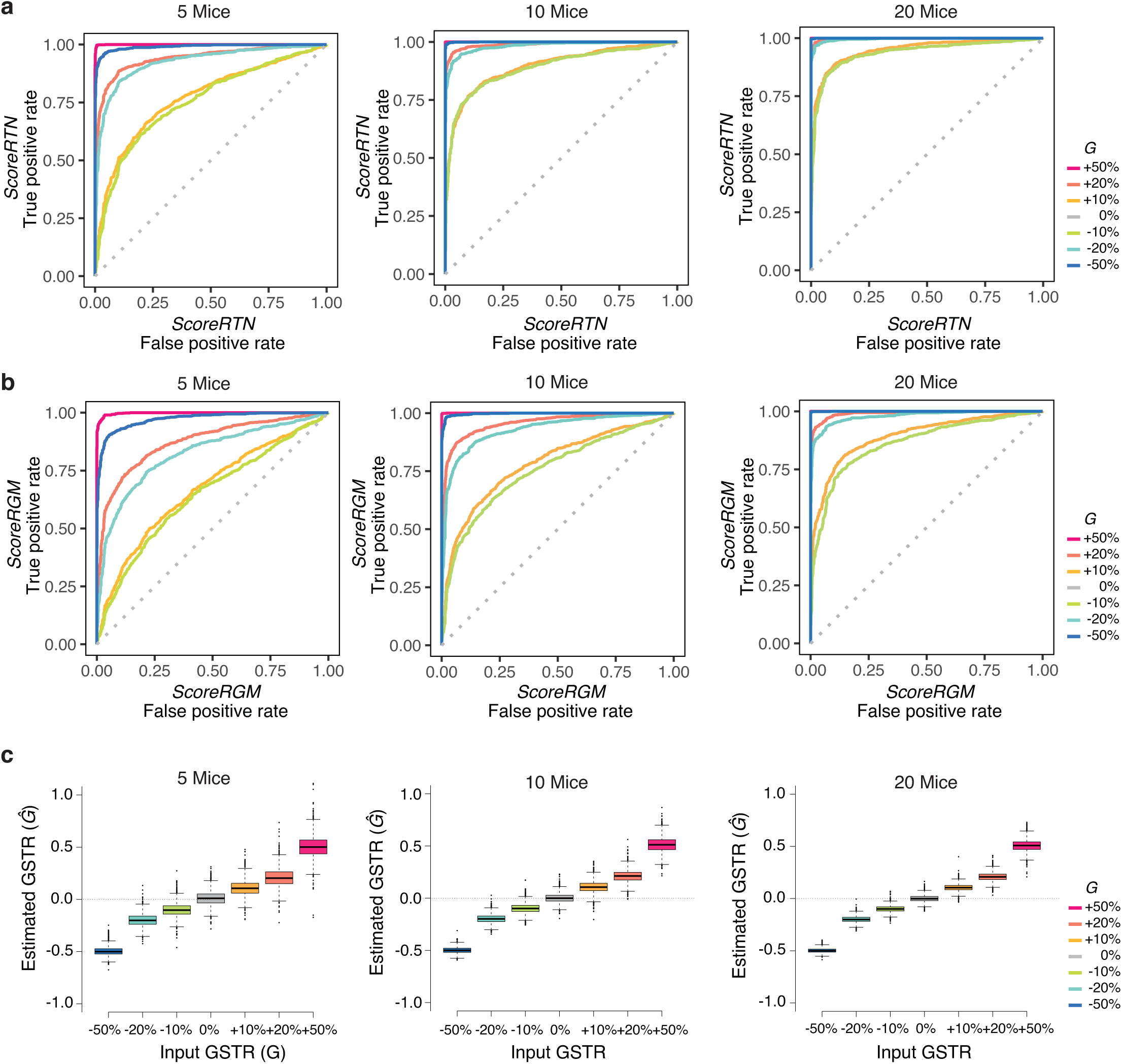
Power analysis for various sample sizes. **a.** The sensitivity and specificity of *ScoreRTN* estimated from simulation of preassigned drug effect (*S*=0.5) and various input GSTR (*G*) when various numbers of mice were used in the treated and untreated group (for example, “10 mice” means that 10 untreated and 10 treated mice were sampled from 8 untreated mice from the pharmacogenomic mapping experiment with replacement for simulation, respectively). The increase in sample size or input GSTR(*G*) leads to a higher power. **b.** The sensitivity and specificity of *ScoreRGM* were estimated with the same parameter setting as in a. **c.** Estimated GSTR (*Ĝ*) by combining *ScoreRTN* and *ScoreRGM* from the above simulations with various input GSTR (*G*). The estimated GSTRs are unbiased and were more accurate with larger sample size.

**Supplemental Figure 5.**
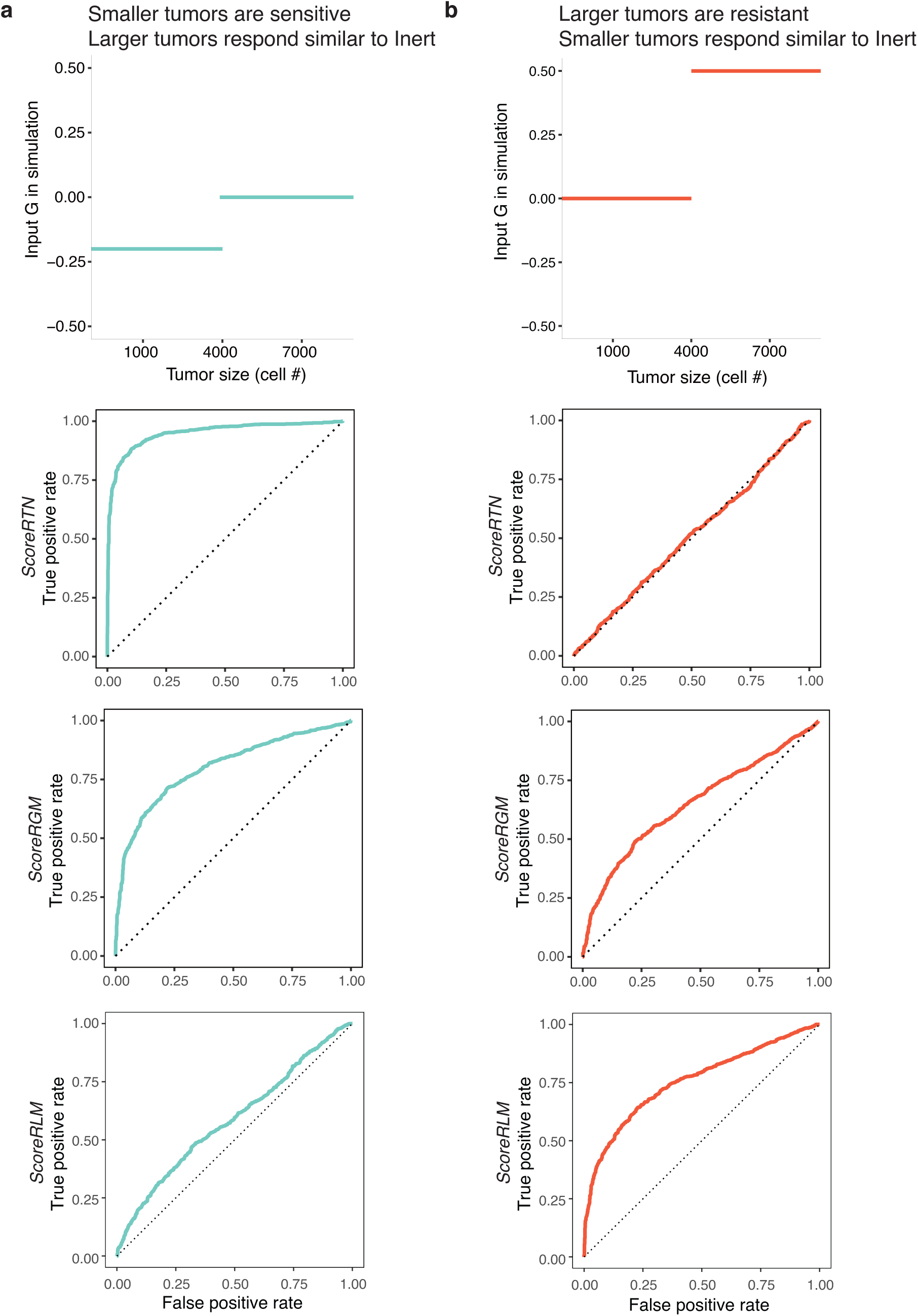
Examples of scenarios where one score outperforms the other. **a.** Five untreated and treated mice were simulated as in Supplementary Fig 4 with *S*=0.5. The top panel shows the preassigned input GSTR (G) on non-Inert tumors, where tumors smaller than 4000 cells showed an additional 20% reduction in size. When small tumors respond more, *ScoreRTN* outperforms *ScoreRGM* in detecting genotype-specific responses. **b.** The simulation was the same as panel a except for the preassigned input GSTR. The top panel shows the input GSTR (*G*), where tumors larger than 4000 cells respond less well to the drug compared with Inert tumors, resulting in tumors being 50% larger than expected when *G*=0. When large tumors respond less, *ScoreRTN* is not a sensitive metric to detect genotype-specific responses; however, *ScoreRGM* retains some power to detect genotype-specific responses. Other summary statistics, such as *ScoreRLM* that compares the relative LN mean for tumors between the treated and untreated mice, can also be used to identify GSTR. Compared with *ScoreRTN* and *ScoreRGM*, *ScoreRLM* has the lowest power when larger tumors respond similar to Inert tumors, and has the highest power when only larger tumors show GSTR.

**Supplemental Figure 6.**
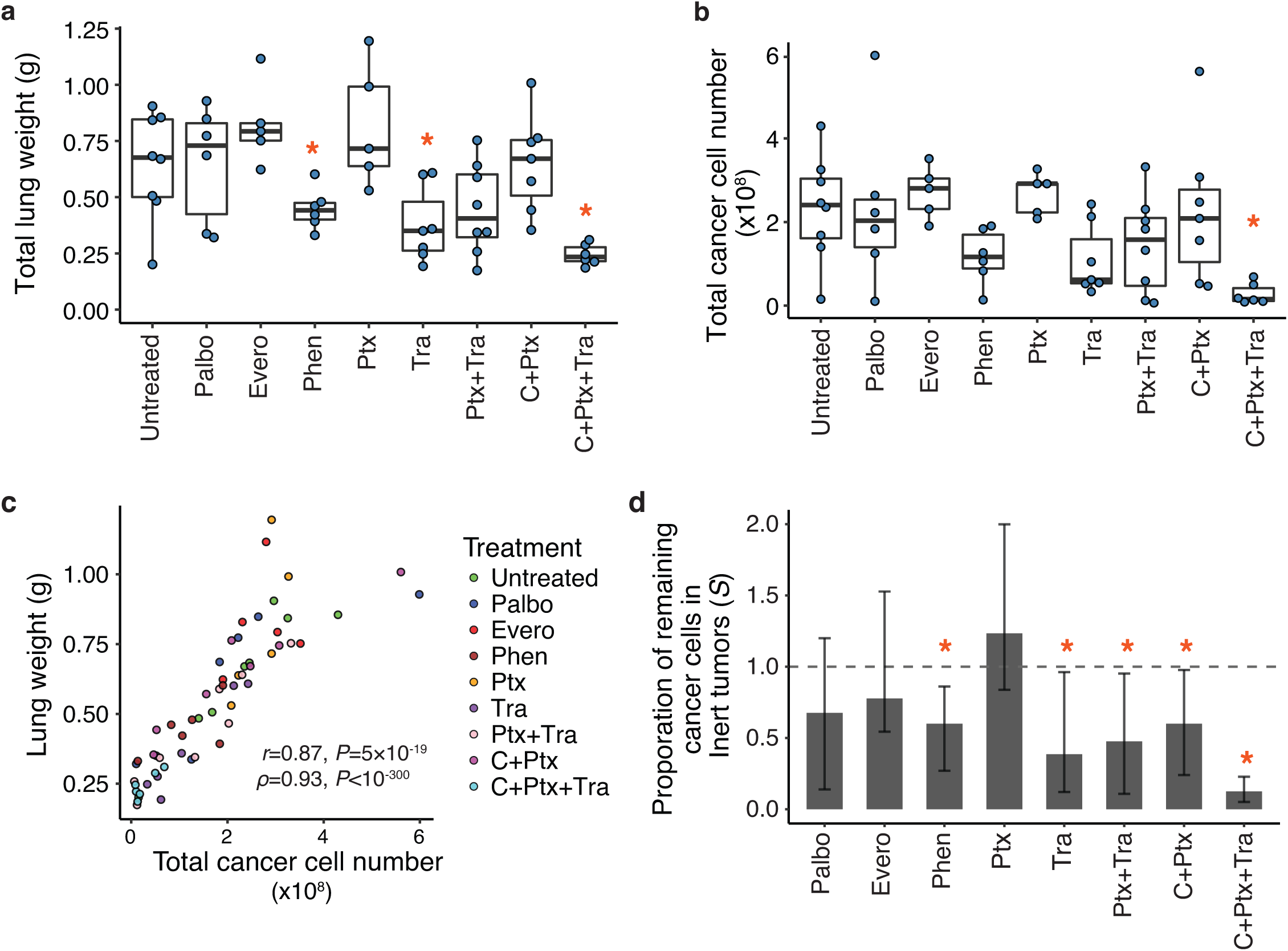
Overall treatment responses of mice to therapies. **a.** Boxplot showing the effects of treatment on mouse lung weight. Each blue dot is the lung weight of a mouse. Several treatments lead to reduced tumor burden sufficient to dramatically reduce total lung weight. Stars below the name of the treatment indicate significant reductions in lung weight by the treatment (*P*<0.05, Mann-Whitney U test). **b.** Boxplot showing the effect of treatment on the total cancer cell number. Each blue dot is the total cancer cell number of a mouse. The total cancer cell number in each lung was determined by converting the number of reads containing a sgID to cell number based on the read number of the three spike-ins with known cell number counts. Stars indicate significant reductions in total cancer cell number by the treatment (*P*<0.05, Mann-Whitney U test). **c.** Total lung weight is highly correlated with the total cancer cell number. **d.** The proportion of remaining cancer cells for Inert tumors *S* for each treatment is estimated by matching the distribution of Inert tumors in the treated and untreated mice. Stars indicate significant reductions in the remaining tumor cells (*P*<0.05 by bootstrap).

**Supplemental Figure 7.**
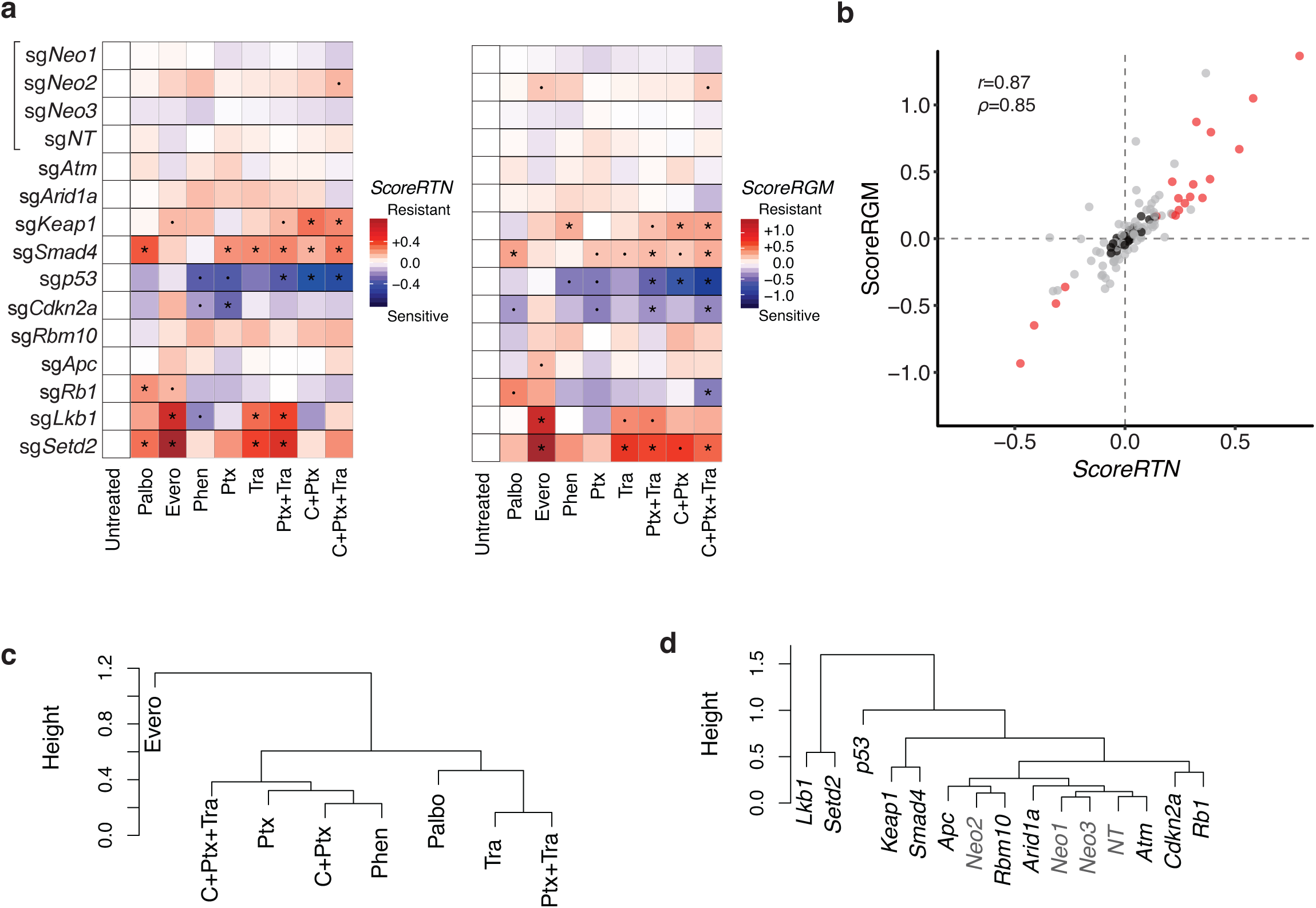
ScoreRTN and ScoreRGM for all genotypes across all treatments and experiments. **a.** Heatmap for *ScoreRTN* and *ScoreRGM* for the pharmacogenomic mapping experiment outlined in Figure 2e. “·” indicates a marginally significant GSTR (*P*<0.1), and “*” indicates a significant GSTR (*P*<0.05). *ScoreRTN* and *ScoreRGM* are integrated to generate *Ĝ* shown in Figure 2f. **b.** *ScoreRTN* and *ScoreRGM* are highly correlated. Red dots show interactions that are significant by one score and at least marginally significant by the other score. Gray dots are the rest of GSTR that doesn’t fit the above criteria. Black dots show all GSTR for Inerts, all with small magnitudes and non-significant p-values. **c.** Hierarchical clustering of the treatments in the pharmacogenomic mapping experiment based on *Ĝ* with complete linkage. Combo treatments clustered close to their corresponding monotherapies. **d.** Hierarchical clustering of the genes in the pharmacogenomic mapping experiment based on *Ĝ* with complete linkage. *Rb1* and *Cdkn2a* are in the same biological pathway and were clustered closely. *Lkb1* and *Setd2* were previously shown to be functionally redundant (Rogers *et al*. 2018) and were clustered closely.

**Supplemental Figure 8.**
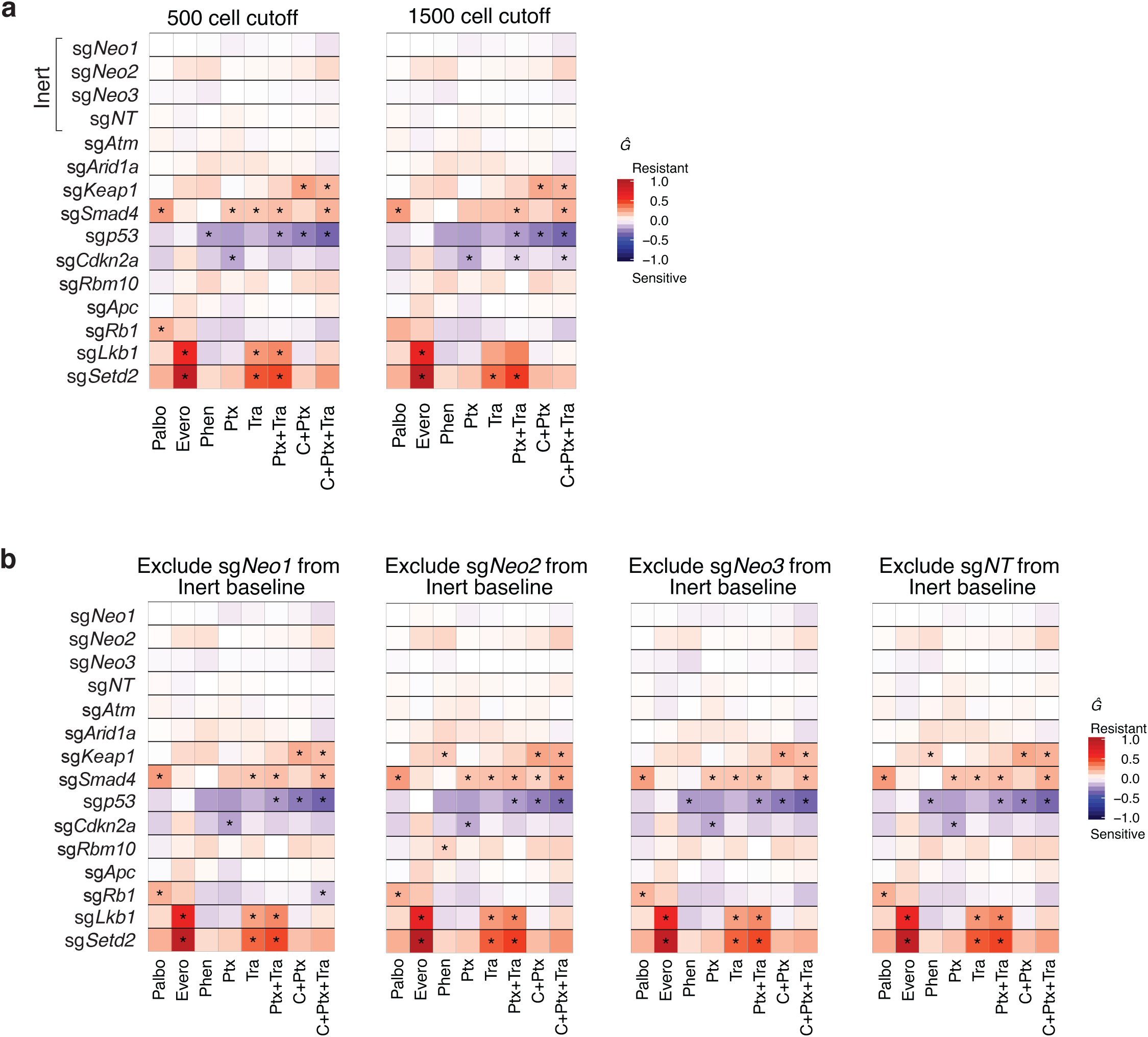
Estimates of GSTRs are robust to changes in minimum tumor size cutoff and the composition of the Inert tumors. **a.** Throughout most analyses in this manuscript, we include tumors in untreated mice that are calculated to have over 1000 cells (a cutoff of 1000 cells). However, using 500 cells or 1500 cells as the tumor size cutoffs generated very similar results. The correlation between *Ĝ* using 500 or 1500 cells as cutoff versus using 1000 cell cutoff was 0.998 and 0.997 for linear correlation, and 0.997 and 0.997 for rank correlation, respectively. Compared to using the original 1000 cell cutoff, 100% and 97.5% of the GSTR identified by Ĝ were in the same direction, respectively. Among 19 significant GSTRs, 18 and 13 significant GSTRs were reidentified using the 500 and 1500 cell cutoff, respectively. Thus, the identified genotype-specific therapeutic responses are almost unaltered by using different cell number cutoffs. **b.** Throughout most analyses in this manuscript, we aggregate tumors with sg*Neo1*, sg*Neo2*, sg*Neo3* and sg*NT* as the Inert tumors. Excluding any of the Inerts yields similar results. The correlation between *Ĝ* leaving out one of the Inerts versus using all four Inerts was over 0.998 for linear correlations and over 0.994 for rank correlations. Over 95.8% of the Significant *Ĝ* results were in the same direction. At least 17 of the 19 significant GSTRs were reidentified in each setting.

**Supplemental Figure 9.**
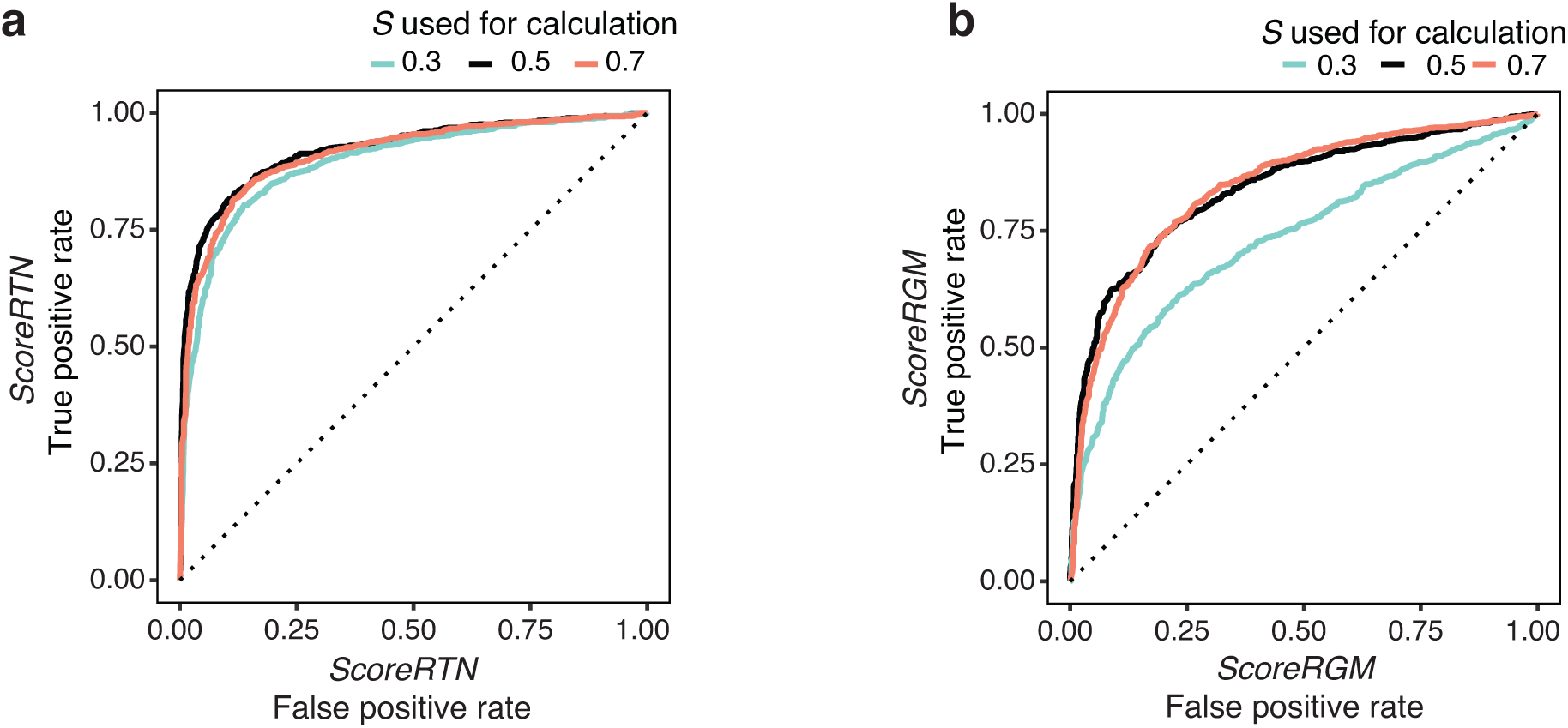
Estimates of GSTR by *ScoreRTN* and *ScoreRGM* are robust to inaccurate estimation of *S*. **a.** The calculation of *ScoreRTN* and *ScoreRGM* both depend on our estimate of the overall drug responses (represented as *S* - the remaining fraction of cancer cells for Inert tumors). In the simulation, we tested the impact of our estimates of *S* being inaccurate. Instead of estimating the effect size based on observed *Inert* tumors, we assign *S* to be a constant value, taking three discrete values 0.3, 0.5, and 0.7 when we know the preassigned drug effect is 0.5. ROC curves were plotted for detecting GSTRs with Input GSTR (*G)*=0.2. Incorrect estimation of *S* only slightly impacts the power of *ScoreRTN*. **b.** Similar simulation is performed for *ScoreRGM* and Incorrect estimation of S only slightly impacts the power of *ScoreRGM*.

**Supplemental Figure 10.**
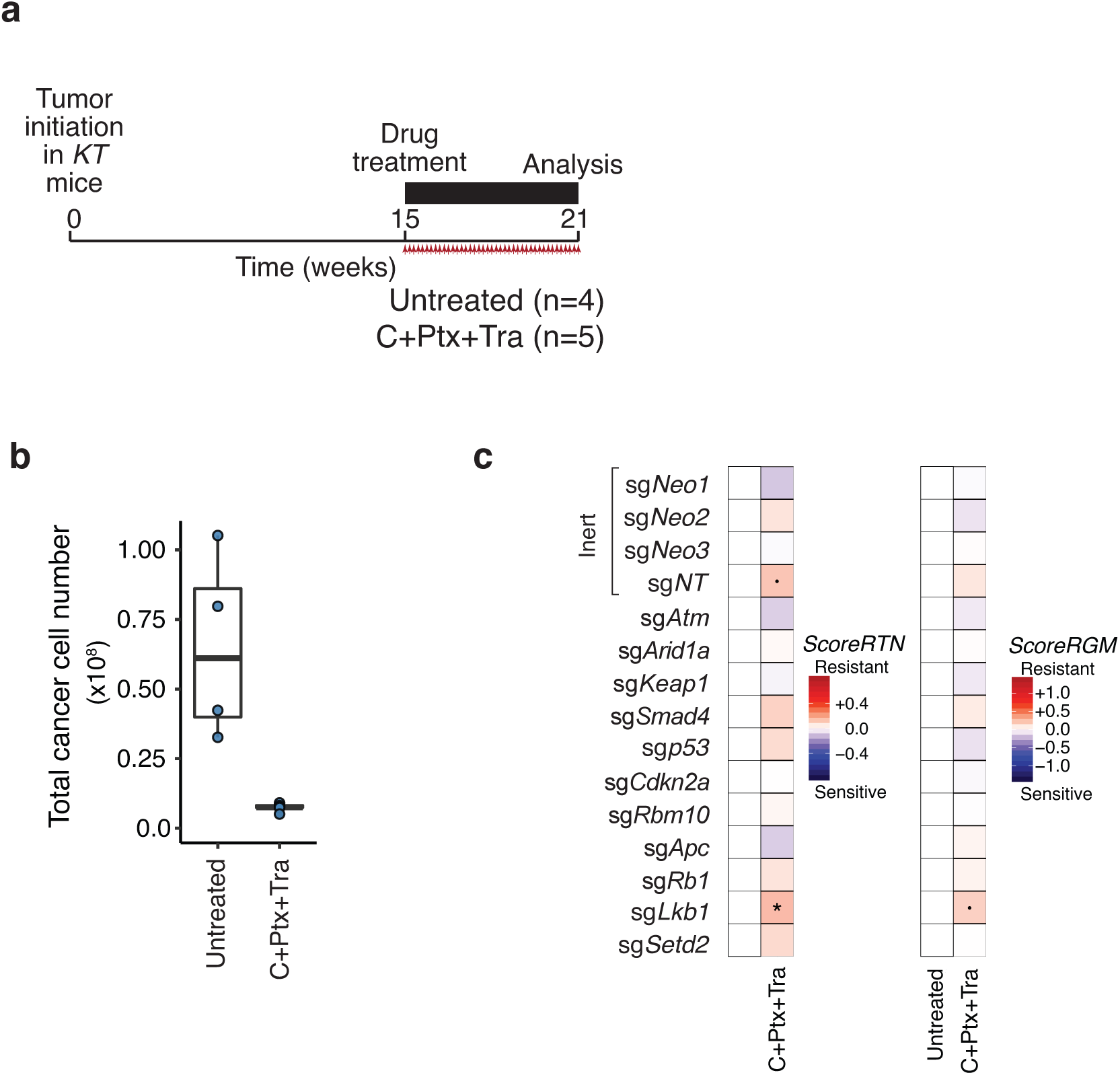
Only 1 weak GSTRs were identified in the negative control experiment in *KT* mice without Cas9. **a.** Outline of a negative control experiment. In this negative control experiment, we initiated tumors in *KT* mice (which lack Cas9) with Lenti-TS^Pool^/Cre followed by a 6-week treatment with Carboplatin+Paclitaxel+Trametinib (C+Ptx+Tra). We chose this treatment, and prolonged treatment time, to generate maximal tumor responses which would most stringently test our analytical models. Given that the mice lack Cas9, all sgRNA are functional Inert, and we expected to find no differences in responses of tumors with different sgRNAs. The number of mice in each group is indicated. **b.** Boxplot showing the effect of treatment on the total cancer cell number. Each blue dot is the total cancer cell number of a mouse. The total cancer cell number in each lung was determined by converting the number of reads containing a sgID to cell number based on the read number of the three spike-ins with known cell number counts. The total cancer cell numbers were significantly different across the two groups. **c.** Heatmap for *ScoreRTN* and *ScoreRGM* for the negative control experiment. “·” indicates a marginally significant GSTR (*P*<0.1), and “*” indicates a significant GSTR (*P*<0.05). Note the relatively low magnitudes of estimated effect relative to those shown in Supplementary Figure 7a. *ScoreRTN* and *ScoreRGM* are integrated to generate *Ĝ* shown in Figure 2g. Apart from tumors with sg*Lkb1* showing significant and marginally significant GSTRs with small magnitudes by *ScoreRTN* and *ScoreRGM*, respectively, other genes were non-significant when comparing the C+Ptx+Tra treated mice to untreated *KT* mice.

**Supplemental Figure 11.**
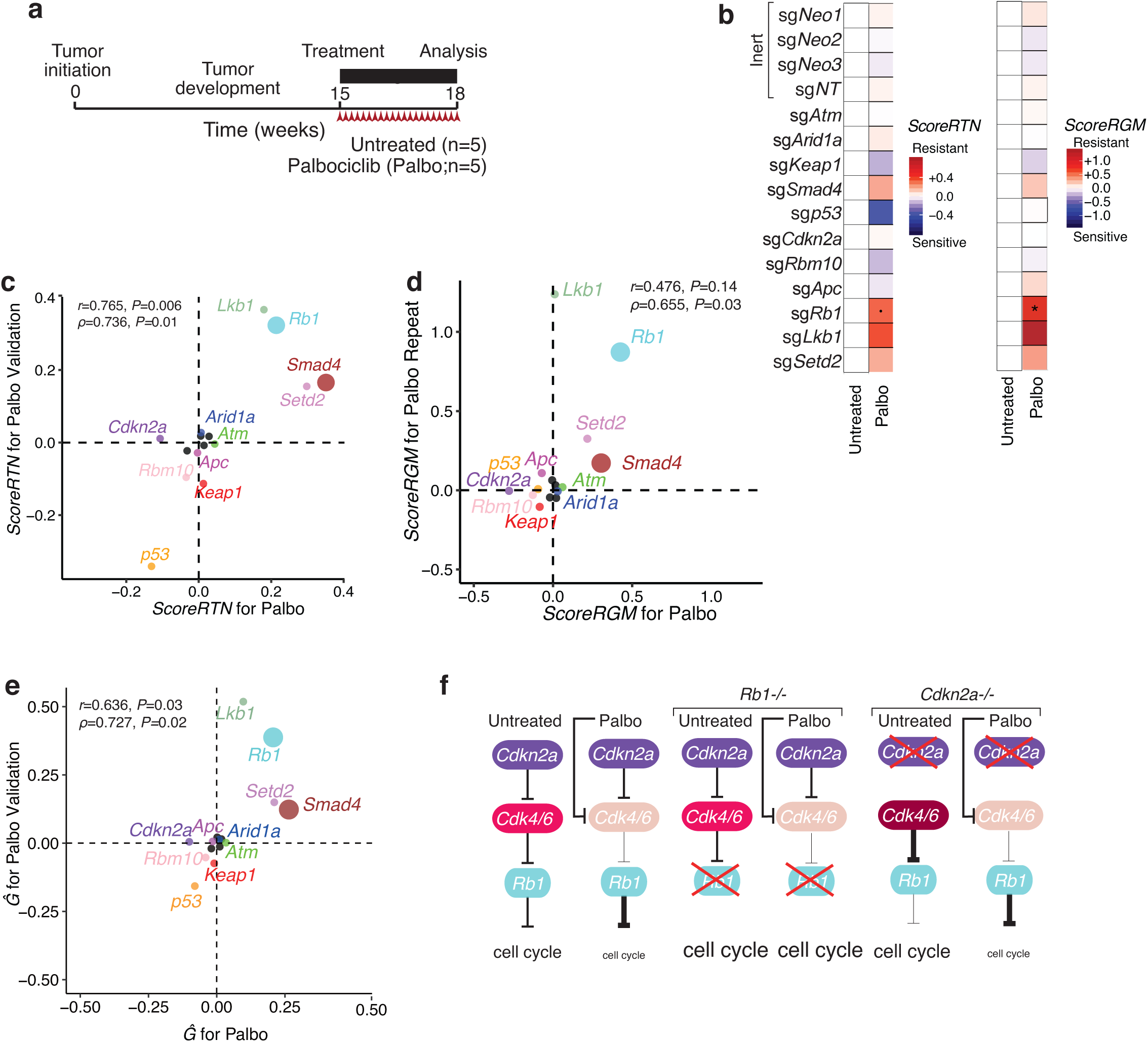
GSTR to Palbociclib identified in a repeat experiment. **a.** Outline of the replicate experiment to identify GSTR to Palbociclib (Palbociclib repeat experiment). **b.** Heatmap for *ScoreRTN* and *ScoreRGM* for the Palbociclib repeat experiment. “·” indicates a marginally significant GSTR (*P*<0.1), and “*” indicates a significant GSTR (*P*<0.05). *ScoreRTN* and *ScoreRGM* are integrated to generate *Ĝ* shown in Figure 2h. **c,d,e**. Across two independent experiments of palbociclib, the identified genotype-specific therapy responses (GSTRs) were recaptured for both *ScoreRTN* (**c**), *ScoreRGM* (**d**), and *Ĝ* (**e**). Large dots indicate significant GSTRs in at least one experiment, and small dots indicate that GSTR was significant in neither experiment. Pearson’s (*r*) and Spearman’s correlations (*ρ*) were calculated. **f**. Palbociclib works as a *Cdk4/6* inhibitor. *Rb1* as the downstream effector is expected to be resistant to Palbociclib. *Cdkn2a* functions as a tumor suppressor upstream in the same pathway; therefore, the growth advantage conferred by the inactivation of *Cdkn2a* will be removed by the Palbociclib treatment, showing drug sensitivity. While *Rb1-*inactivation lead to significant resistance in both experiments, *Cdkn2a*-inactivation lead to marginally significant sensitivity for both *ScoreRTN* and *ScoreRGM* in the pharmacogenomic mapping experiment. Darker color intensities of Cdk4/6 represents higher activity.

**Supplemental Figure 12.**
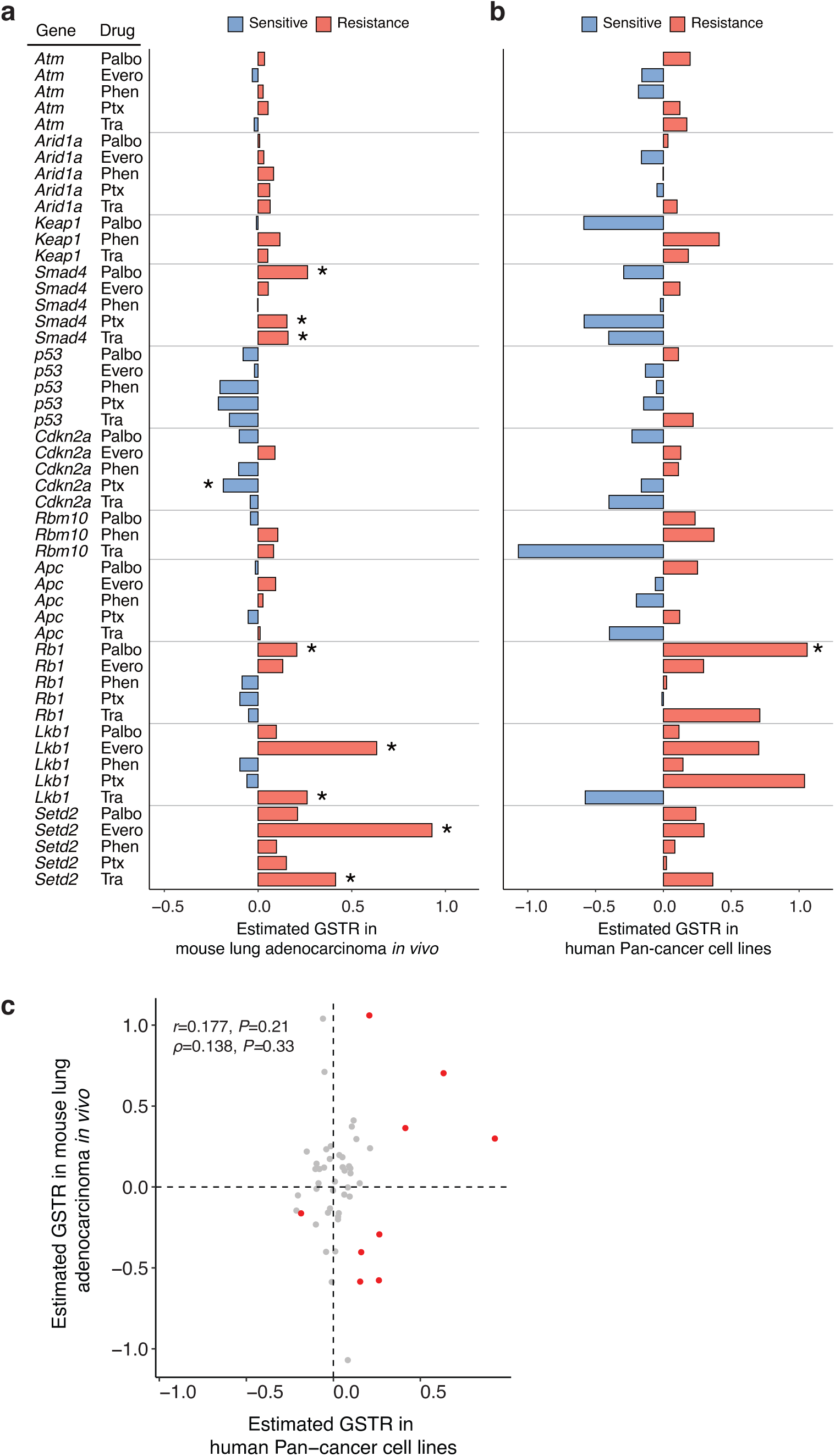
Comparison with the GDSC human cell line database. **a.** GSTR estimated from our study from mouse lung tumors *in vivo*. **b.** GSTR reported by GDSC for human PAN cancer cell lines. Genes are in the same order as a. **c.** Correlation between GSTR estimated in our study and the cancer cell line study. The significant GSTRs in our study are highlighted in red.

**Supplementary Figure 13.**
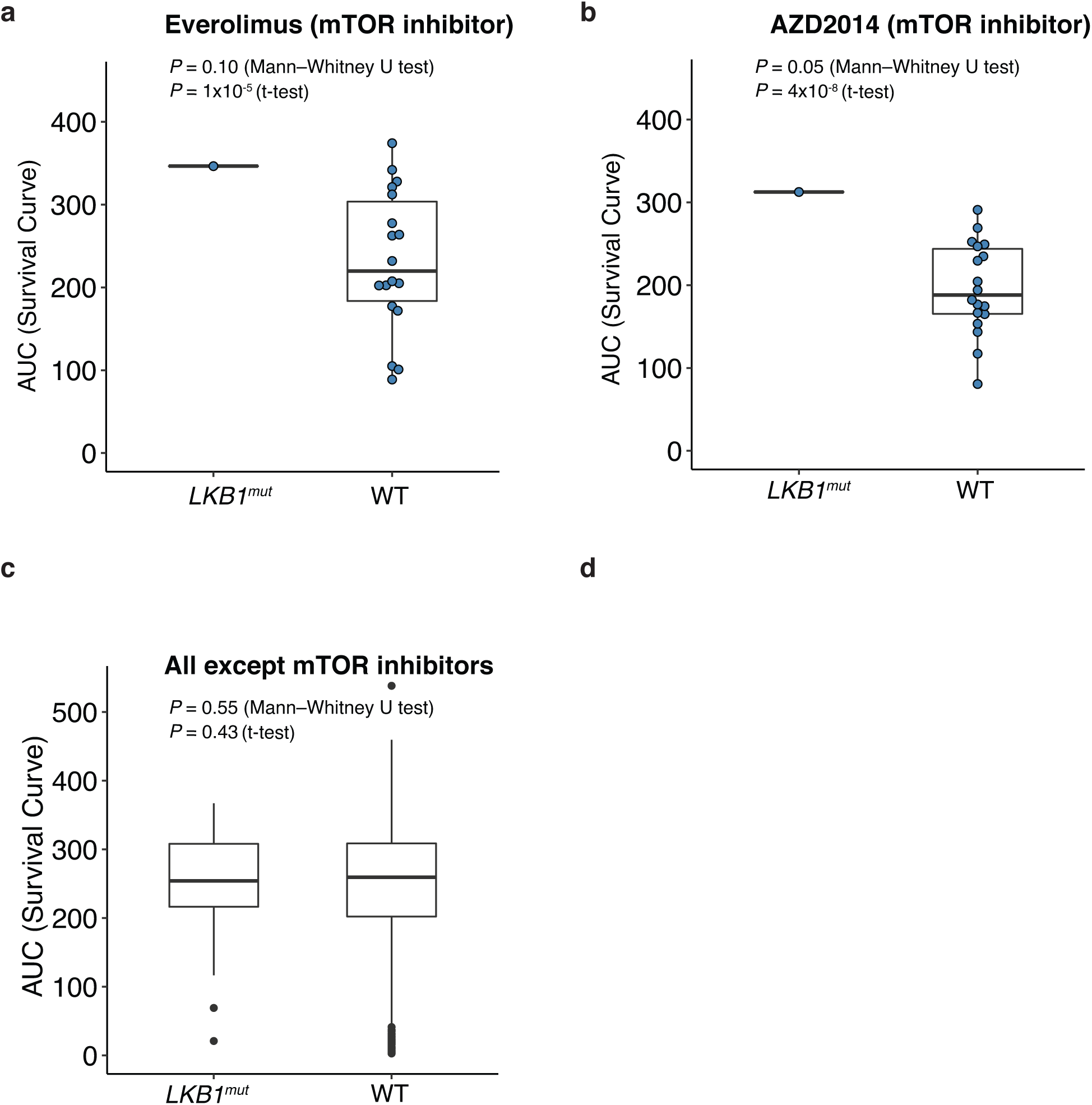
Correlation of *LKB1*-mutation with resistance to mTOR inhibition in patient-derived cancer cells from patients with lung adenocarcinoma. **a,b.** Jin-Ku Lee *et al.* quantified the area under the curve (AUC) of the dose-response curves as a proxy for the pharmacological responses of patient-derived cancer cells from 462 patients to 60 treatments (the Cancer-Drug eXplorer database). We focused on the 21 primary cell cultures derived from patients with lung adenocarcinomas. Among them, one had an LKB1 mutation and those cells showed resistance to two mTOR inhibitor, everolimus (a), and AZD2014 (b). *P*-values were calculated from one-sided Mann-Whitney U-test or one-sided t-test. **c.** The *LKB1-*mutant tumor-derived cells do not show general resistance to treatments. Another explanation for the resistance of that one sample to mTOR inhibition could be that those cancer cells (with the *LKB1* mutation) are just more resistant to therapies in general. The AUC for *LKB1* to all treatments other than mTOR inhibitors were not different from cell lines that are wild-type in *LKB1*. 985 data points were used for comparison and we did not plot individual data points for better visualization.

**Supplementary Figure 14.**
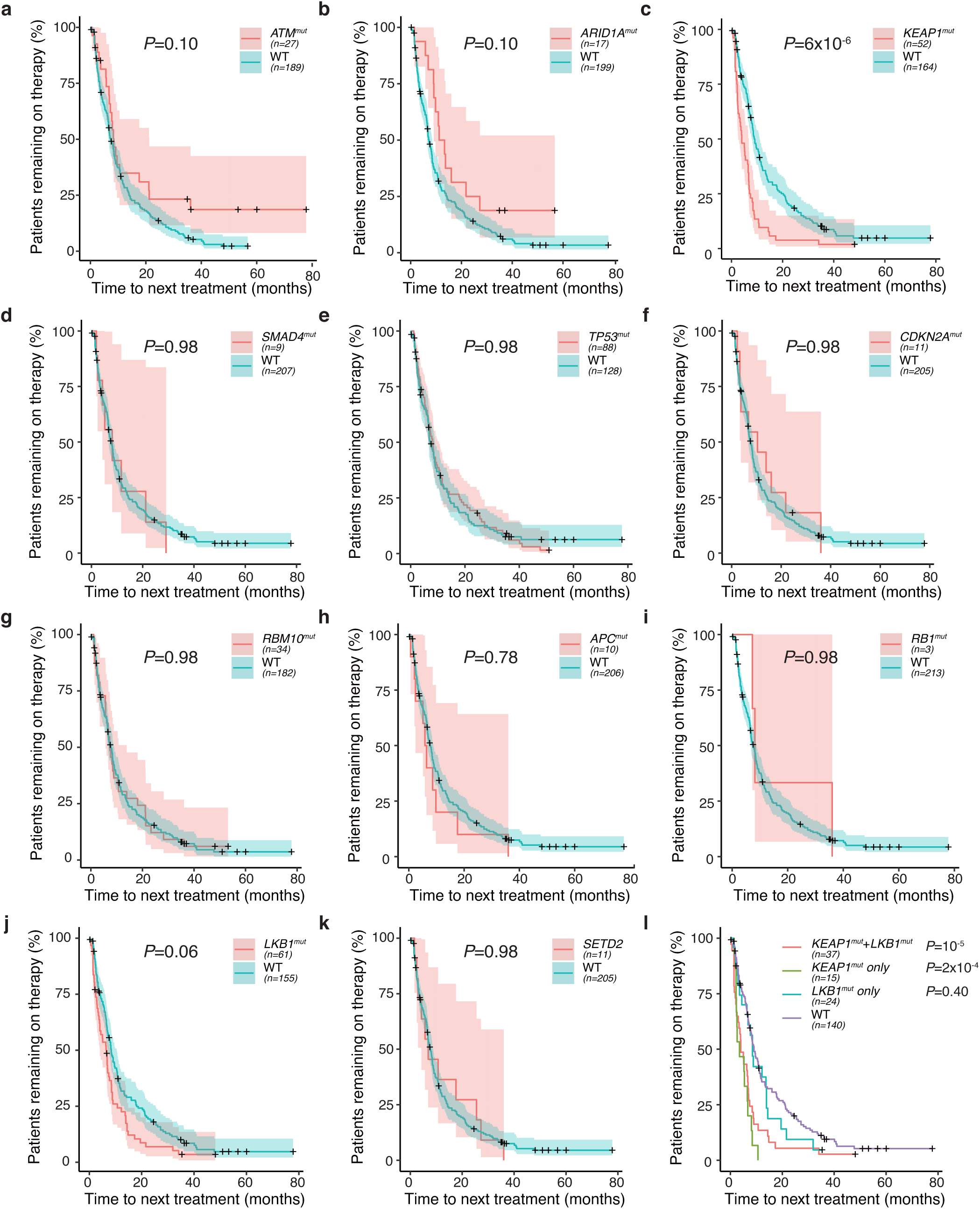
Patient response to chemotherapy represented by Kaplan–Meier estimator plot. **a-k.** Kaplan-Meier curve (with 95% confidence interval in shading) of time-to-next-treatment (months) for patients with or without mutation of the gene of interest. Longer time-to-next-treatment is associated with better responses to chemotherapy. Here, we analyzed data from a total of 216 patients with mutations in KRAS codon 12, 13, or 16. *P*-values were calculated from the Mantel-Haenszel test with FDR correction. Genotypes compared are indicated for each plot and the number of patients in each group is shown. **l**. *KEAP1* mutations commonly co-occur with *LKB1* mutations. We divide patients into four groups based on their mutation status of *LKB1* and *KEAP1*, and found that the nominal resistance of the *LKB1* mutant group (see Figure S13j) seems to be due to the effect of co-incident *KEAP1* mutations. Importantly, the absence of resistance conferred by LKB1 mutation is consistent with our mouse model data. *P*-values were calculated relative to WT using the Mantel-Haenszel test with FDR correction.

**Supplementary Figure 15.**
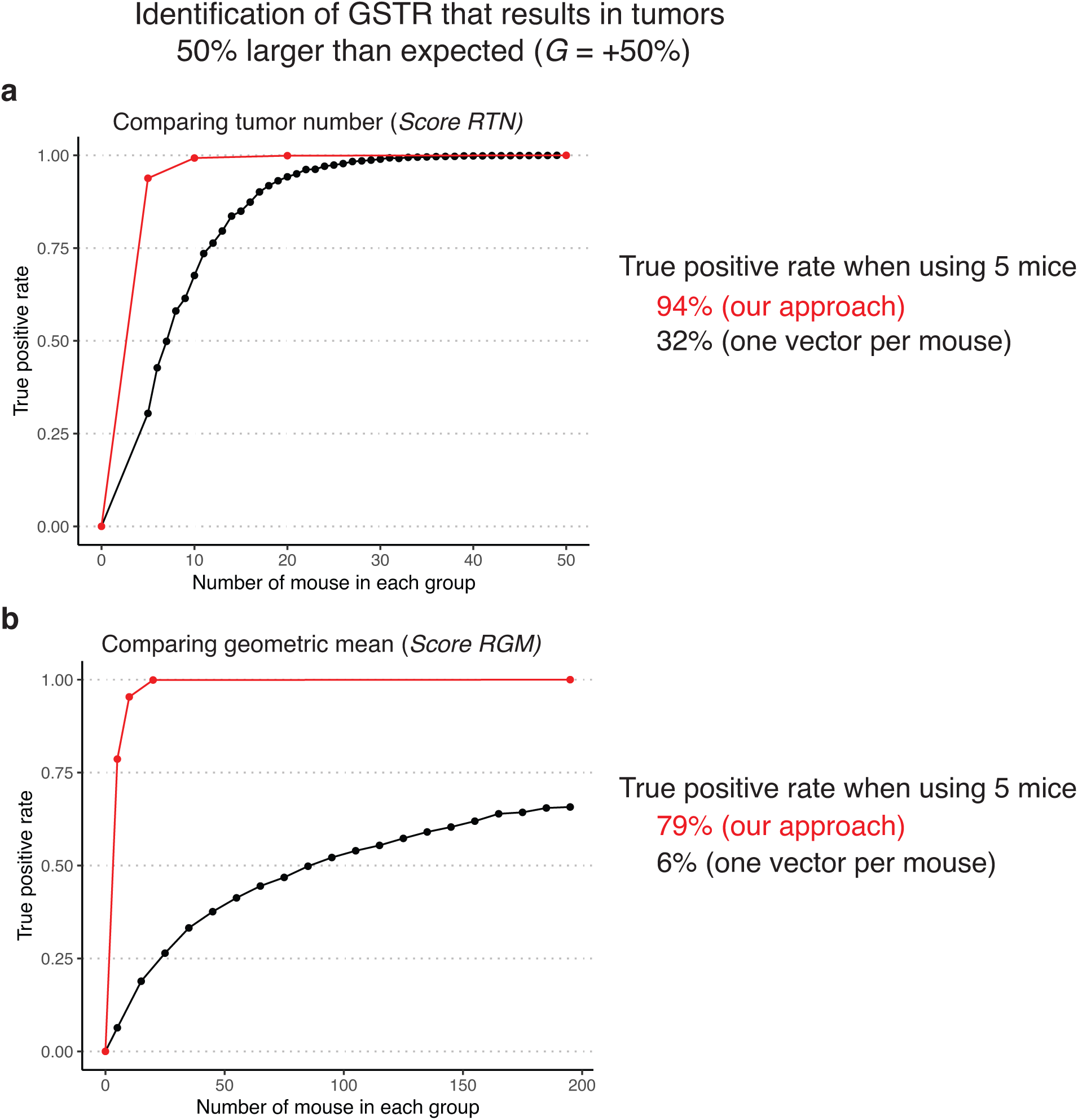
The Tuba-seq platform greatly reduces the amount of work and costs for identifying genotype-specific therapeutic responses. **a.** The sensitivity of using our Tuba-seq method (*ScoreRTN* metric) with multiplexed vectors is higher than having one vector per mouse and comparing the number (and/or size) of tumors between untreated and treated cohorts. We performed 1000 simulations for each sample size and calculated the probability of identifying the preassigned GSTR. In our study, using five mice in each group has a high true positive rate of 94% for identifying GSTRs with *G*=+50%. For the one vector per mouse approach, to achieve only a 50% true positive rate, we would need ∼7 mice per vector, and a total of 672 mice (7 mice for each of the 12 genotypes under eight treatments). The cost of doing the study with one genotype per mouse would be 17-times the cost to achieve half of our true positive rate. Keep in mind that this includes a 17-fold increase in the cost of animal housing, 17-fold increase in viral vector production, 17-fold increase in the amount of each drug need, 17-fold increase in the number of mouse dosings, and 17-fold increase in the number of Tuba-seq libraries and sequencing costs. Labor-related costs would also be much higher, and the total cost of a one genotype per mouse study would exceed several million dollars. Following the same logic, to achieve a similar true positive rate of 94%, ∼50 times more mice would be needed with proportional increases in all costs. **b.** Sensitivity for using our Tuba-seq method (*ScoreRGM* metric) with multiplexed vectors is much higher than having one vector per mouse and comparing the geometric mean of tumors between the untreated and treated mice. Note that assuming uniform GSTR, the metric that compares the tumor numbers has higher statistical power than the metric comparing the geometric means, and the x-axes are plotted on different scales for better visualization. Using the size metric *ScoreRGM* alone, having five mice in each group results in a true positive rate of 79% in identifying GSTR with *G*=+50%. For the one vector per mouse approach, to achieve a 50% true positive rate, we would need ∼85 mice per vector, which is a total of 8160 mice for performing the pharmacogenomic mapping experiment. The total cost for a study like that would be astronomical.

