## Supplementary Table 1 for "Quantitative *in vivo* analyses reveal a complex pharmacogenomic landscape in lung adenocarcinoma"

**Table S1. Genotype-specific treatment responses**

| sgID | Treatment | ScoreGSTR[1] | Significance[2] | *ScoreRTN*[3] | *pRTN*[4] | *ScoreRGM*[5] | *pRGM*[6] |
|---|---|---|---|---|---|---|---|
| Atm | Palbo | 0.034 | | 0.044 | 0.295 | 0.060 | 0.274 |
| Arid1a | Palbo | 0.009 | | 0.007 | 0.423 | 0.030 | 0.437 |
| Keap1 | Palbo | -0.009 | | 0.013 | 0.385 | -0.084 | 0.368 |
| Smad4 | Palbo | 0.264 | * | 0.351 | 0.003 | 0.304 | 0.030 |
| p53 | Palbo | -0.080 | | -0.130 | 0.145 | -0.094 | 0.363 |
| Cdkn2a | Palbo | -0.101 | | -0.107 | 0.126 | -0.278 | 0.090 |
| Rbm10 | Palbo | -0.041 | | -0.035 | 0.573 | -0.127 | 0.243 |
| Apc | Palbo | -0.015 | | -0.003 | 0.388 | -0.071 | 0.274 |
| Rb1 | Palbo | 0.207 | * | 0.214 | 0.029 | 0.425 | 0.072 |
| Lkb1 | Palbo | 0.097 | | 0.180 | 0.151 | 0.011 | 0.509 |
| Setd2 | Palbo | 0.211 | | 0.298 | 0.022 | 0.217 | 0.243 |
| Atm | Evero | -0.031 | | -0.038 | 0.349 | -0.064 | 0.325 |
| Arid1a | Evero | 0.031 | | 0.044 | 0.442 | 0.045 | 0.474 |
| Keap1 | Evero | 0.094 | | 0.125 | 0.099 | 0.141 | 0.209 |
| Smad4 | Evero | 0.054 | | 0.077 | 0.172 | 0.076 | 0.289 |
| p53 | Evero | -0.019 | | -0.033 | 0.463 | -0.014 | 0.525 |
| Cdkn2a | Evero | 0.090 | | 0.134 | 0.103 | 0.101 | 0.307 |
| Rbm10 | Evero | 0.048 | | 0.046 | 0.394 | 0.127 | 0.381 |
| Apc | Evero | 0.094 | | 0.097 | 0.142 | 0.215 | 0.095 |
| Rb1 | Evero | 0.132 | | 0.136 | 0.100 | 0.294 | 0.158 |
| Lkb1 | Evero | 0.633 | * | 0.581 | 0.000 | 1.049 | 0.002 |
| Setd2 | Evero | 0.928 | * | 0.792 | 0.000 | 1.366 | 0.000 |
| Atm | Phen | 0.027 | | 0.039 | 0.310 | 0.037 | 0.375 |
| Arid1a | Phen | 0.083 | | 0.119 | 0.187 | 0.105 | 0.257 |
| Keap1 | Phen | 0.116 | | 0.120 | 0.137 | 0.263 | 0.050 |
| Smad4 | Phen | -0.002 | | -0.014 | 0.413 | 0.029 | 0.459 |
| p53 | Phen | -0.203 | | -0.306 | 0.072 | -0.387 | 0.075 |
| Cdkn2a | Phen | -0.103 | | -0.154 | 0.074 | -0.166 | 0.196 |
| Rbm10 | Phen | 0.106 | | 0.134 | 0.109 | 0.176 | 0.193 |
| Apc | Phen | 0.027 | | 0.042 | 0.299 | 0.029 | 0.436 |
| Rb1 | Phen | -0.085 | | -0.101 | 0.142 | -0.203 | 0.148 |
| Lkb1 | Phen | -0.097 | | -0.200 | 0.076 | -0.002 | 0.457 |
| Setd2 | Phen | 0.098 | | 0.051 | 0.419 | 0.362 | 0.183 |
| Atm | Ptx | 0.053 | | 0.075 | 0.189 | 0.074 | 0.270 |
| Arid1a | Ptx | 0.062 | | 0.087 | 0.224 | 0.088 | 0.283 |
| Keap1 | Ptx | -0.014 | | -0.028 | 0.391 | 0.000 | 0.511 |
| Smad4 | Ptx | 0.154 | * | 0.216 | 0.026 | 0.182 | 0.076 |
| p53 | Ptx | -0.212 | | -0.327 | 0.059 | -0.392 | 0.090 |
| Cdkn2a | Ptx | -0.186 | * | -0.272 | 0.013 | -0.362 | 0.053 |

| | | | | | | | |
|---|---|---|---|---|---|---|---|
| Rbm10 | Ptx | 0.038 | | 0.073 | 0.206 | 0.004 | 0.415 |
| Apc | Ptx | -0.053 | | -0.065 | 0.188 | -0.115 | 0.188 |
| Rb1 | Ptx | -0.097 | | -0.102 | 0.143 | -0.270 | 0.134 |
| Lkb1 | Ptx | -0.060 | | -0.041 | 0.427 | -0.218 | 0.256 |
| Setd2 | Ptx | 0.151 | | 0.209 | 0.108 | 0.187 | 0.176 |
| Atm | Tra | -0.021 | | -0.024 | 0.331 | -0.046 | 0.380 |
| Arid1a | Tra | 0.064 | | 0.101 | 0.239 | 0.059 | 0.353 |
| Keap1 | Tra | 0.052 | | 0.070 | 0.218 | 0.081 | 0.306 |
| Smad4 | Tra | 0.160 | * | 0.229 | 0.016 | 0.173 | 0.087 |
| p53 | Tra | -0.153 | | -0.230 | 0.142 | -0.266 | 0.214 |
| Cdkn2a | Tra | -0.042 | | -0.027 | 0.383 | -0.155 | 0.256 |
| Rbm10 | Tra | 0.083 | | 0.126 | 0.173 | 0.087 | 0.306 |
| Apc | Tra | 0.011 | | 0.011 | 0.475 | 0.030 | 0.402 |
| Rb1 | Tra | -0.051 | | -0.046 | 0.305 | -0.155 | 0.222 |
| Lkb1 | Tra | 0.262 | * | 0.309 | 0.023 | 0.406 | 0.089 |
| Setd2 | Tra | 0.414 | * | 0.389 | 0.006 | 0.796 | 0.015 |
| Atm | Ptx+Tra | 0.011 | | 0.016 | 0.452 | 0.015 | 0.455 |
| Arid1a | Ptx+Tra | 0.039 | | 0.072 | 0.294 | 0.013 | 0.427 |
| Keap1 | Ptx+Tra | 0.108 | | 0.130 | 0.085 | 0.198 | 0.093 |
| Smad4 | Ptx+Tra | 0.178 | * | 0.244 | 0.010 | 0.213 | 0.050 |
| p53 | Ptx+Tra | -0.221 | * | -0.314 | 0.035 | -0.485 | 0.016 |
| Cdkn2a | Ptx+Tra | -0.096 | | -0.089 | 0.188 | -0.296 | 0.043 |
| Rbm10 | Ptx+Tra | 0.007 | | 0.041 | 0.346 | -0.073 | 0.313 |
| Apc | Ptx+Tra | -0.004 | | 0.003 | 0.418 | -0.030 | 0.441 |
| Rb1 | Ptx+Tra | -0.023 | | -0.001 | 0.483 | -0.119 | 0.251 |
| Lkb1 | Ptx+Tra | 0.321 | * | 0.385 | 0.008 | 0.445 | 0.082 |
| Setd2 | Ptx+Tra | 0.473 | * | 0.518 | 0.003 | 0.669 | 0.018 |
| Atm | C+Ptx | 0.027 | | 0.018 | 0.540 | 0.092 | 0.227 |
| Arid1a | C+Ptx | 0.026 | | 0.055 | 0.211 | -0.014 | 0.509 |
| Keap1 | C+Ptx | 0.231 | * | 0.295 | 0.003 | 0.312 | 0.031 |
| Smad4 | C+Ptx | 0.108 | * | 0.142 | 0.046 | 0.167 | 0.089 |
| p53 | C+Ptx | -0.281 | * | -0.413 | 0.010 | -0.649 | 0.005 |
| Cdkn2a | C+Ptx | -0.057 | | -0.053 | 0.237 | -0.171 | 0.103 |
| Rbm10 | C+Ptx | 0.105 | | 0.115 | 0.124 | 0.221 | 0.142 |
| Apc | C+Ptx | 0.017 | | -0.004 | 0.455 | 0.101 | 0.315 |
| Rb1 | C+Ptx | -0.028 | | -0.046 | 0.302 | -0.028 | 0.374 |
| Lkb1 | C+Ptx | -0.037 | | -0.163 | 0.157 | 0.237 | 0.324 |
| Setd2 | C+Ptx | 0.175 | | 0.049 | 0.444 | 0.727 | 0.053 |
| Atm | C+Ptx+Tra | -0.014 | | -0.016 | 0.359 | -0.033 | 0.380 |
| Arid1a | C+Ptx+Tra | -0.057 | | -0.051 | 0.350 | -0.174 | 0.178 |
| Keap1 | C+Ptx+Tra | 0.196 | * | 0.242 | 0.015 | 0.302 | 0.039 |
| Smad4 | C+Ptx+Tra | 0.205 | * | 0.271 | 0.013 | 0.265 | 0.044 |

| | | | | | | | |
|---|---|---|---|---|---|---|---|
| p53 | C+Ptx+Tra | -0.340 | * | -0.475 | 0.009 | -0.933 | 0.001 |
| Cdkn2a | C+Ptx+Tra | -0.093 | | -0.068 | 0.184 | -0.337 | 0.024 |
| Rbm10 | C+Ptx+Tra | 0.097 | | 0.134 | 0.161 | 0.133 | 0.231 |
| Apc | C+Ptx+Tra | 0.049 | | 0.059 | 0.343 | 0.096 | 0.270 |
| Rb1 | C+Ptx+Tra | -0.110 | | -0.091 | 0.130 | -0.374 | 0.030 |
| Lkb1 | C+Ptx+Tra | 0.085 | | 0.064 | 0.334 | 0.265 | 0.149 |
| Setd2 | C+Ptx+Tra | 0.244 | | 0.224 | 0.101 | 0.560 | 0.016 |
| Atm | KT_Ctrl | -0.039 | | -0.063 | 0.342 | -0.042 | 0.331 |
| Arid1a | KT_Ctrl | 0.006 | | 0.009 | 0.351 | 0.008 | 0.445 |
| Keap1 | KT_Ctrl | -0.016 | | -0.014 | 0.596 | -0.047 | 0.410 |
| Smad4 | KT_Ctrl | 0.049 | | 0.077 | 0.109 | 0.050 | 0.254 |
| p53 | KT_Ctrl | 0.018 | | 0.057 | 0.419 | -0.060 | 0.291 |
| Cdkn2a | KT_Ctrl | -0.001 | | 0.002 | 0.397 | -0.014 | 0.519 |
| Rbm10 | KT_Ctrl | 0.006 | | 0.014 | 0.415 | -0.006 | 0.577 |
| Apc | KT_Ctrl | -0.027 | | -0.064 | 0.307 | 0.027 | 0.407 |
| Rb1 | KT_Ctrl | 0.027 | | 0.041 | 0.332 | 0.030 | 0.399 |
| Lkb1 | KT_Ctrl | 0.093 | * | 0.125 | 0.028 | 0.138 | 0.068 |
| Setd2 | KT_Ctrl | 0.032 | | 0.061 | 0.315 | 0.002 | 0.460 |
| Atm | Palbo_Rep | 0.002 | | -0.004 | 0.527 | 0.019 | 0.503 |
| Arid1a | Palbo_Rep | 0.013 | | 0.028 | 0.483 | -0.006 | 0.515 |
| Keap1 | Palbo_Rep | -0.074 | | -0.113 | 0.189 | -0.105 | 0.361 |
| Smad4 | Palbo_Rep | 0.123 | | 0.166 | 0.173 | 0.172 | 0.206 |
| p53 | Palbo_Rep | -0.157 | | -0.341 | 0.293 | 0.007 | 0.502 |
| Cdkn2a | Palbo_Rep | 0.005 | | 0.011 | 0.502 | -0.004 | 0.500 |
| Rbm10 | Palbo_Rep | -0.053 | | -0.096 | 0.274 | -0.030 | 0.455 |
| Apc | Palbo_Rep | 0.006 | | -0.028 | 0.404 | 0.108 | 0.341 |
| Rb1 | Palbo_Rep | 0.387 | * | 0.324 | 0.092 | 0.872 | 0.050 |
| Lkb1 | Palbo_Rep | 0.518 | | 0.366 | 0.111 | 1.237 | 0.110 |
| Setd2 | Palbo_Rep | 0.150 | | 0.155 | 0.371 | 0.325 | 0.519 |

[1] *ScoreGSTR* – Combined GSTR score calculated from *ScoreRTN* and *ScoreRGM*.

[2] *Significance* – Only cases that were significant in one score (*P*<0.05) and at least marginally significant in the other score (*P*<0.1) were reported as significant in the combined score.

[3] *ScoreRTN* – Genotype-specific drug response score calculated from relative tumor number.

[4] *pRTN* – The *P*-value calculated for *ScoreRTN* by bootstrap.

[5] *ScoreRGM* – Genotype-specific drug response score calculated from relative geometric mean.

[6] *pRGM* – The *P*-value calculated for *ScoreRGM* by bootstrap.
